# Chromatin reprogramming and transcriptional regulation orchestrate embryogenesis in hexaploid wheat

**DOI:** 10.1101/2022.01.21.477188

**Authors:** Long Zhao, Xuelei Lin, Yiman Yang, Xiaomin Bie, Hao Zhang, Jinchao Chen, Xuemei Liu, Hao Wang, Jiafu Jiang, Xiangdong Fu, Xiansheng Zhang, Jun Xiao

**Author notes:** These authors contributed equally to this work (Xuelei Lin, Yiman Yang).

## Abstract

Embryogenesis represents the beginning of life cycle, but our understanding of the regulatory circuitry in plants is far lagged to animals. Here, we draw a transcriptome trajectory and chromatin landscape profile during embryogenesis of most cultivated crop hexaploid wheat, highlighting large-scale chromatin reconfiguration and distinct proximal and distal transcriptional regulation in defining cell fate transition. Upon fertilization, H3K27ac and H3K4me3 resetting were correlated with maternal genome silence, while *de novo* building of chromatin accessibility activated zygotic genome. Global depletion of H3K27me3 in pre-embryo results in a permissive chromatin environment with gain-of-chromatin accessibility, allowing subsequent hierarchical cis- and trans-regulation network mediated by key factors, such as LEC1, MYB, ZHD, LEC2, governing embryo pattern formation. By contrast, H3K27me3 restoration coordinating with chromatin compaction in developmental genes attenuated totipotency and prohibited extensive organogenesis during embryo maturation. In addition, dynamic biased expression of homeolog triads and diverse expression profiles after polyploidization were observed. This is correlated with asymmetric transposon elements insertion in accessible proximal and distal regions. Thus, our study revealed a plant-specific chromatin reprogramming process in facilitating the hierarchical transcription regulation circuits mediated “inverse hourglass model” and unveiled epigenetic regulation of evolutionary divergence among different sub-genome in shaping embryogenesis in polyploidy wheat.

## Introduction

Embryogenesis, during which fusion of parental terminally differentiated gametes following fertilization generates an entirely new organism, represents the beginning of development and ensures the continuation of the life cycle for both plants and animals^1–4^. Plant embryogenesis can be generally divided into three steps: early-embryogenesis, during which a totipotent zygote was formed following fertilization; mid-embryogenesis, during which the major cell lineages and embryo body pattern was built; and late-embryogenesis, during which nutrition was accumulated in mature embryo followed by seed dormancy^1, 2^. Comparisons between animals and plants’ embryonic development highlight both conservation and difference. Animals and plants share a similar cell fate determination process-cell division followed by cell identity specification during embryonic development. In addition, both the transcriptional patterns of animals and plants go through the same transcriptomic hourglass model where a conserved phylotypic period during mid-embryogenesis punctuating divergent early- and late-stage between species with a phylum and the inverse hourglass model where gene expression patterns during early- and late-stage are more comparable than mid-stage^5, 6^. Nevertheless, there are also differences between the two kingdoms: maternal dominant versus both parental contribution to early-embryogenesis, completed versus partial organogenesis, with dormancy or not, etc^1, 7, 8^. Besides the morphological and transcriptome similarity and difference, a large gap is the chromatin dynamics comparison between two kingdoms, due to the lack of comprehensive epigenomic study in plants.

Epigenetics enables one genome to give rise to a spectrally and phenotypically diverse collection of cells and tissues in response to developmental and environmental cues^9, 10^. The technological advances for epigenetics landscape profiling, such as histone modifications, DNA accessibility and DNA methylation, etc., greatly facilitate the discovery of cis-regulatory elements and other non-coding genomic features associated with development, environment, disease and cancer^11–13^. Epigenetic regulatory logic ensures the precise progress and timing of maternal-to-zygotic transition (MZT), zygotic gene activation (ZGA), cell-fate decision, and lineage-specific differentiation during animal embryogenesis^14, 15^. A large-scale epigenetic remodeling occurred during embryogenesis, including histone modifications, chromatin accessibility and 3D chromatin organization, as well as DNA methylation^4, 16^. H3K4me3 is rapidly depleted after fertilization but re-established during major ZGA and its active removal is important for ZGA initiation^17^. Besides, a noncanonical form of H3K4me3 was present in both humans and mice at distal intergenic regions, though the exact function is not clear^18^. Global erasure of H3K27me3 occurred during early embryogenesis, but with parental genomic allelic difference manner^4^. As well, H3K27me3 could be inter-generationally inherited from the maternal genome to regulator enhancer activity during MZT^19^. Plus, H3K4me3 and H3K27me3 co-occupy lineage-specific genes at the blastocyst stage suggest a critical role of bivalency in poised state maintenance and gene expression regulation^20^. Major chromatin accessibility reorganization is critical for epigenetic reprogramming to convert terminally differentiated gametes into a totipotent state^21^. Open chromatin was progressively established during early embryogenesis and showed a major increase in 8-cell embryos in mouse^22, 23^, correlated with gene activation. However, previous studies on plant embryogenesis have mainly focused on the transcriptional change and functional identity of specific genes at certain embryo development stages^8, 24–28^. A systematic view of how epigenomic landscape contributes to genes regulation in cell fate transition, lineage specification and cellular metabolic accumulation during plant embryogenesis is lacking, especially for crops^29^.

Wheat, as the most cultivated crop worldwide, provides 20% of the calories and protein daily consumed and is important for human sustention. The majority of global wheat production comes from two species, the hexaploid bread wheat (*Triticum aestivum*), which accounts for 95% of global wheat production and tetraploid pasta wheat (*Triticum turgidum* var *durum*). Hexaploid wheat (AABBDD) is the result of two times hybridization and polyploidization events of three putative diploid wild grass progenitor species, *Triticum monococcum* (AA), an *Aegilops speltoides*-related species (BB), and *Aegilops tauschii* (DD)^30, 31^. A recent finding investigated the transcriptional landscape of polyploidy wheat and diploid ancestors during embryogenesis^27^. Through comparison among different ploidy wheat, the evolutionary divergence of gene expression and contribution of A, B, D subgenome to grain development in polyploidy wheat were characterized^27^. Yet, without further information such as epigenomic profiles, it would be hard to dissect the complex transcription regulation network governing such programmed cell fate transition process and elucidation of driving force for the ontogenetic divergence along with the evolution of polyploidy wheat species. Besides, the elucidation of transcriptional complexity and epigenome atlas during embryonic development could be helpful to study in other developmental processes and plant breeding applications, such as parthenogenesis and somatic embryogenesis^32, 33^.

Here, we investigate various histone modifications and chromatin accessibility of eight-stage samples across embryonic development of hexaploid wheat, providing a systematic view of transcriptome trajectory and chromatin landscape profiles. Thousands of distal genomic regulatory elements and putative enhancer-gene linkages are identified to drive embryogenesis. A plant-specific large-scale chromatin state reconfiguration and transcriptional regulation circuitry were characterized during embryonic phase transition. Further, we provide evidence for an explanation of the regulatory mechanisms underlining evolutionary divergence among different sub-genomes and the ‘inverse hourglass model’ of transcription variability during hexaploid wheat embryogenesis. This data thus provides a comprehensive resource of genetic and epigenomic annotations for researchers studying transcriptional regulation, embryonic development and polyploid evolution.

## Result

### Charting the chromatin landscapes during wheat embryogenesis

Despite the importance of transcription (re)programming and epigenetic regulation during plant embryogenesis, a comprehensive survey is limited for both model plants and crops^29^. This is partly due to the difficulty to get robust epigenomic data from a limited sample of developing embryos. Here, the Cleavage Under Targets and Tagmentation (CUT&Tag) and Assay for Transposase Accessible Chromatin with high-throughput sequencing (ATAC-seq) were used to tackle this challenge^34, 35^. In a pilot run, CUT&Tag showed not only high correlation with published Chromatin immunoprecipitation (ChIP)-seq data (correlation >0.8) but also high repeatability and sensitivity with lower nuclei input (∼1,000 nuclei/reaction) and sequencing depth (∼1/3-1/20 of ChIP-seq) (Supplementary Fig. 1a-c). ATAC-seq from about 5,000 nuclei showed logical peak numbers, good signal-to-noise ratio, reasonable fragment size and genomic distribution, and fully meet the demand of transcription factors (TFs) footprint calling (Supplementary Fig. 1d-h).

With this capability, the ‘reference epigenome’ of eight embryonic developmental stages were generated (DPA 0, 2, 4, 6, 8, 12, 16, 22, days post anthesis), including histone modifications, histone variant and RNA Polymerase II profile and chromatin accessibility, as well as transcriptome (Fig. 1a). In general, various chromatin marks showed featured distribution patterns and logical correlation with transcription (Supplementary Fig. 2a, b). As well, pair-wise correlations analysis showed that active markers (H3K27ac, H3K9ac, H3K4me3, Pol II, H3K36me3, H3K4me1, H2A.Z and ATAC) correlate positively with one another but showed varying degrees of exclusivity with repressive markers (H3K27me3, H3K9me2, H3K9me3) (Supplementary Fig. 2c). Unsupervised PCA and hierarchical clustering of RNA-seq and ATAC-seq revealed a continuous trajectory which can be divided into five major groups roughly corresponding to fertilization, pre-embryo, transition, differentiation and maturation of embryo (Fig. 1b, e). In contrast, core histone modifications showed a phase transition pattern, which revealed a global reconfiguration of chromatin at a certain stage (Supplementary Fig. 2d). GO enrichment of time-lapse correlated modules revealed that cell cycle and DNA replication-related genes were activated during early embryogenesis, organ and tissue specification-related processes were distinguished during mid-development, and nutrient accumulation and dormancy were built during the late stage (Fig. 1c). This is evident in the evolutionary conservation of monocots and dicots embryogenesis^27, 28^. In particular, some known factors involved in embryo pattern formation, such as *WOX11*, *LEC2, ARR6* and *ABI5*, showed similar expression patterns during embryogenesis among wheat, *Brachypodium* and *Arabidopsis* but not the other genes, such as *YUC1* (Fig. 1d). Furthermore, chromatin accessibility was progressively built until DPA 8, transition stage for embryo patterning, whereas gradually closed during differentiation stage (Fig. 1f). Notably, the majority of ATAC-seq peaks are more than 3Kb even 20Kb away from transcriptional start site (TSS) of genes (Fig. 1g), which is different from plants of small genome size^34, 36^. Interestingly, more ATACseq peaks distributed even further away from TSS in B subgenome as compared to A or D subgenome (Fig. 1g), coincidently with the fact that B subgenome is larger than A or D^31^.

**Fig. 1.**
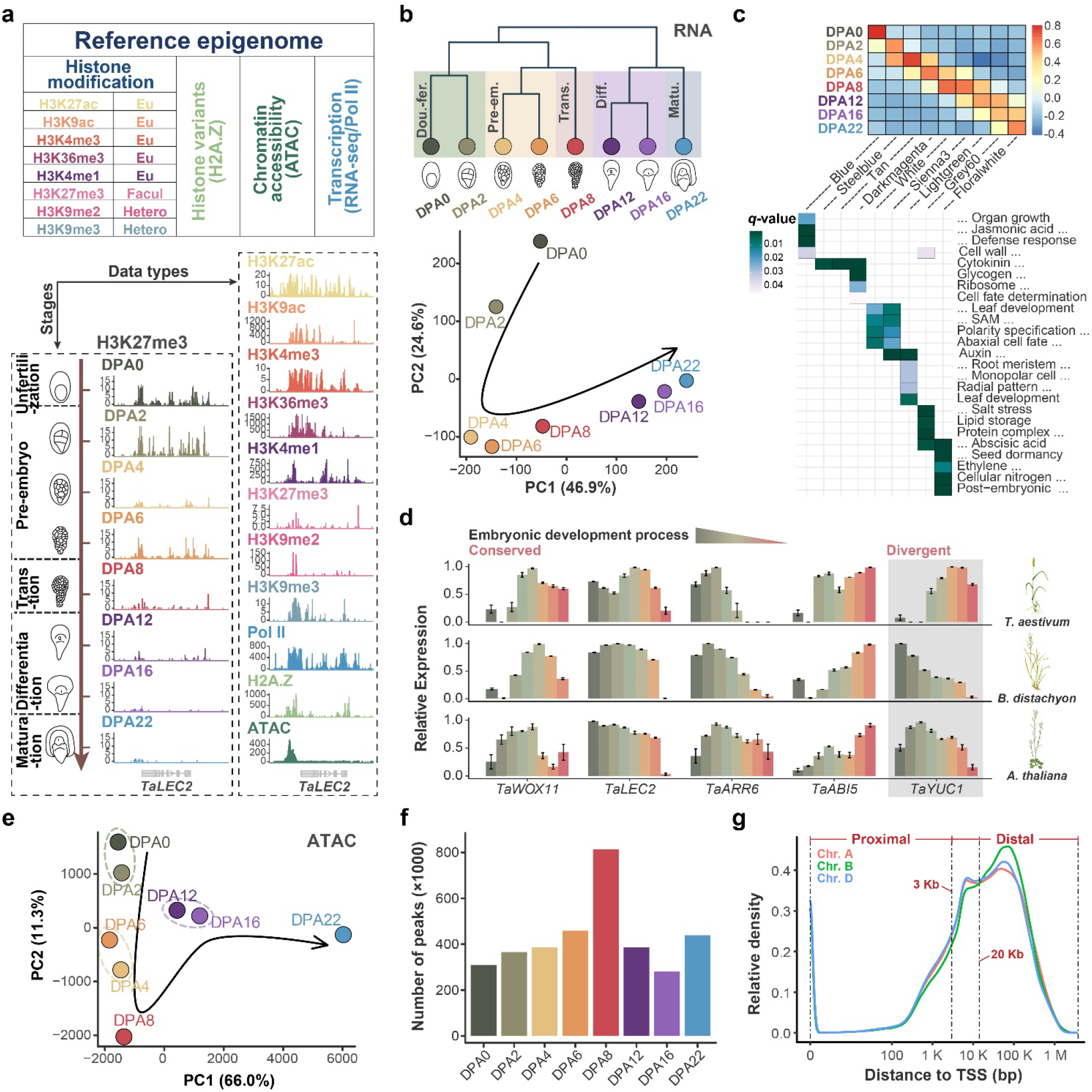
Charting chromatin landscapes of wheat embryogenesis. **a**, Experimental design and major axes of the data series: data types and sampling developmental stages, DPA: days post anthesis. **b**, Cluster dendrogram and PCA of transcriptome showing five distinct development stages: around double fertilization (DPA 0/2), pre-embryo (DPA 4/6), transition (DPA 8), differentiation (DPA 12/16), and maturation (DPA 22). **c**, Representative modules from WGCNA cluster and enrichment of corresponding GO. **d**, The conserved and diversified expression pattern of representative genes between monocot (*T. aestivum* and *B. distachyon*) and dicot (*A. thaliana*) during embryogenesis. **e-g,** Dynamic pattern of genome-wide chromatin accessibility measured by ATACseq during embryogenesis. PCA analysis (**e**), peak numbers dynamics (**f**), and peak distribution in different sub-genomes (**g**).

Though it is hard to isolate purified embryo at early embryonic development stages (DPA 0, 2, 4), the progressive change of transcription and chromatin accessibility presented here indicates a logical and meaningful cellular process. Thus, our ‘reference epigenome’ could capture the dynamics of transcription and potential transcriptional regulation features during wheat embryogenesis process.

### Distinct feature of proximal and distal chromatin accessible regions and correlation with transcription regulation

Accessible chromatin can expose regulatory DNA sequences to transcription factors, and thus emerges as an accurate proxy for transcription regulatory potential and development^37^. In this study, we identified 1,315,547 accessible chromatin regions (ACRs) from ATAC-seq in total, which could be further categorized into genic (g), promoter (p) and distal (d)ACRs based on the location relative to genes, with a better correlation between dACRs, pACRs and gene expression than gACRs (Fig. 2a). Interestingly, the gain and loss of pACRs and dACRs varied largely during embryogenesis (Fig. 2b and Supplementary Fig. 3a, b). Lots of genes gained pACRs after fertilization at DPA2 but dramatically lost during late embryogenesis at DPA1 and DPA16. Whereas dACRs, accounting for about 75% of total ACRs, showed a sharp ‘burst’ at DPA8, the transition stage between cell division and differentiation, then quickly declined at DPA12 (Fig. 2b). Genome-wide ACRs distribution showed that centromere region and distal arms of chromosome including telomere are the ‘hot spot’ for chromatin loose-tight regions, with highly overlapped with Gypsy and CACTA types of TEs. In addition, there was a highly synchronized distribution pattern between ACRs and H3K9me3, but not other histone modifications, at those chromatin eruptions hot spots (Fig. 2b and Supplementary Fig. 3c). This may be correlated with transient expression of TEs just like the activation of retrotransposons in mouse and human 2-cell stage embryo which is critical for the activation of ZGA genes during early embryogenesis^4^. Besides the temporal dynamics, subgenomes of hexaploid wheat showed slight differences for the chromatin accessibility, especially at centromere and telomere regions in addition to the low-colinear chromosome regions (Fig. 2d and Supplementary Fig. 3d).

**Fig. 2.**
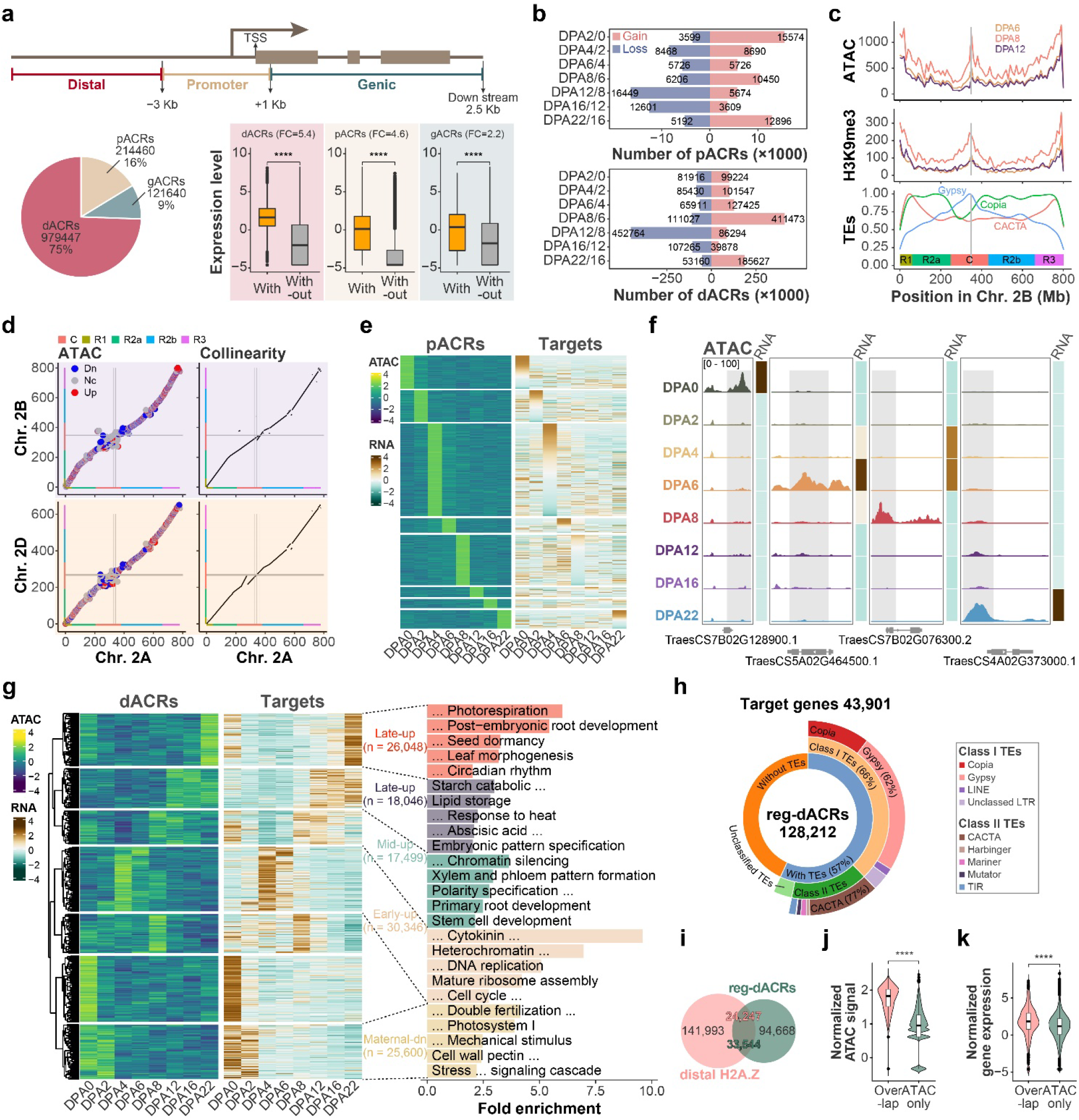
Distinct feature of different ACRs in transcriptional regulation in wheat. **a**, Subdividing ACRs based on distribution pattern around genes, and correlation between different types of ACRs and genes expression. **b**, Distinct gain and loss of pACRs and dACRs between adjacent embryonic developmental stages. **c**, ATAC peaks, H3K9me3 and various types of TEs distributions on chromosome 2B at different embryogenesis stages. **d**, Comparisons of chromatin accessibility of collinear regions among subgenomes, chromosome 2B compared to 2A (up-left) and chromosome 2D with 2A (down-left). Collinear regions were identified based on sequence conservation (right). Color bars on x and y axis indicate the chromosome segments defined by **Consortium (IWGSC) et al., 2018. Grey lines indicate the centromere** for individual chromosomes. **e**, Stage-specific pACRs and corresponding genes expression. **f**, Representative genes in **e.** **g**, Heatmap showing correlation between reg-dACRs and corresponding genes (left) and GO enrichment for different genes sets that were clustered based on the linking reg-dACRs dynamic (right). **h**, Fraction of reg-dACRs overlapped with various types of TEs. **i**, Overlap between reg-dACRs and distal H2A.Z histone variant. **j and k,** Comparisons of ATAC signal intensity (**j**) **and target**s **expression (k) between** reg-dACRs overlapped with H2A.Z and those not overlapped. Mann-Whitney U test (two-sided) was used for **a**, **j** and **k**. * : p <= 0.05; ** : p <= 0.01; ***: p <= 0.001; ****: p <= 0.0001.

The different remodeling pattern of pACR and dACR suggests a distinct regulation weight of gene expression during embryogenesis. Indeed, the stage-abundant pACRs did not perfectly fit corresponding genes, as well as functional pathway enrichment (Fig. 2e, f and Supplementary Fig. 4a). We thus assigned dACRs to corresponding genes using a correlation strategy as the previous report (Method; Supplementary Fig. 4b)^38^. In total, we identified 128,212 regulatory dACR (termed reg-dCAR) linked to 43,901 genes, which accounts for 60% of expressed genes during embryo development and shows higher expression variation (Fig. 2g and Supplementary Fig. 4c). High interaction frequency between reg-dACRs and targets measured by Hi-C and conservation among various species emphasized the accuracy of this distal assignment strategy (Supplementary Fig. 4d, e). Only 35% reg-dACRs are assigned to the nearest genes (Supplementary Fig. 4f, g). Most reg-dACRs have only one target, while one gene could be regulated by multiple dACRs (Supplementary Fig. 4h, i). Genes with few reg-dACRs regulations were enriched for essential biological processes, whereas multiple reg-dACR regulated genes are preferred in stress response and organ development (Supplementary Fig. 4j). The assigned target genes of stage-specific reg-dACRs are well associated with specified developmental stages during embryogenesis (Fig. 2g). Of interest, about 60% of the reg-dACRs were located within TEs of different types (Fig. 2h). Interestingly, the accessibility of reg-dACRs and expression of linked genes were sensitive to TEs classes where they are located (Supplementary Fig. 4k, l). Besides, reg-ACRs were highly overlapped with H2A.Z, a histone variant with looser DNA wrapping as compared to canonical H2A^39^ (Fig. 2i). Indeed, more than 80% H2A.Z signal located within reg-ACRs, and reg-dACRs with H2A.Z showed higher ‘enhancer activity’ with more accessibility and higher expression of targeted genes (Fig. 2i).

Thus, proximal and distal chromatin accessibility undergoes distinct reprogramming patterns during the process of embryogenesis based on genomic features underlying, with different functional output. Meanwhile, a large proportion of long-range distal regulation and asymmetry of chromatin accessibility dynamics among subgenomes exhibit uniqueness for the huge allohexaploid wheat genome.

### Chromatin reprogramming during wheat embryogenesis

To generally classify chromatin states, we integrated eight histone modification marks and Pol II profiles from four embryonic developmental stages (DPA 0, 2, 4, 8) using ChromHMM^40, 41^. Totally 15 chromatin states were categorized, covering 0.02% to 3.5% of the genome respectively (exclude No-signal states) (Fig. 3a). They further grouped into five major functional classes: Promoter (Pr), Enhancer-like (EnL), Transcriptional (Tr), Polycomb group (PcG), and Heterochromatin (Hc) (Fig. 3a). Promoter and enhancer-like classes are enriched for active histone modifications such as H3K27/9ac, H3K4me1/3, chromatin accessibility and histone variant H2A.Z (Fig. 3a). As expected, chromatin states and histone modifications underlying were dramatically altered across developmental stages, with the enhancer-like class varied the most than promoters, just like the case in mouse fetal development^42^ (Fig. 3b, c). This is correlated with the dramatic transcriptome fluctuation (Fig. 1b, c) and regulatory role of dACRs and pACRs (Fig. 2e, h). In addition, the chromatin state pattern at DPA0 is distinguished from other stages (Fig. 3d), revealing a drastic histone modification reprogramming after fertilization.

**Fig. 3.**
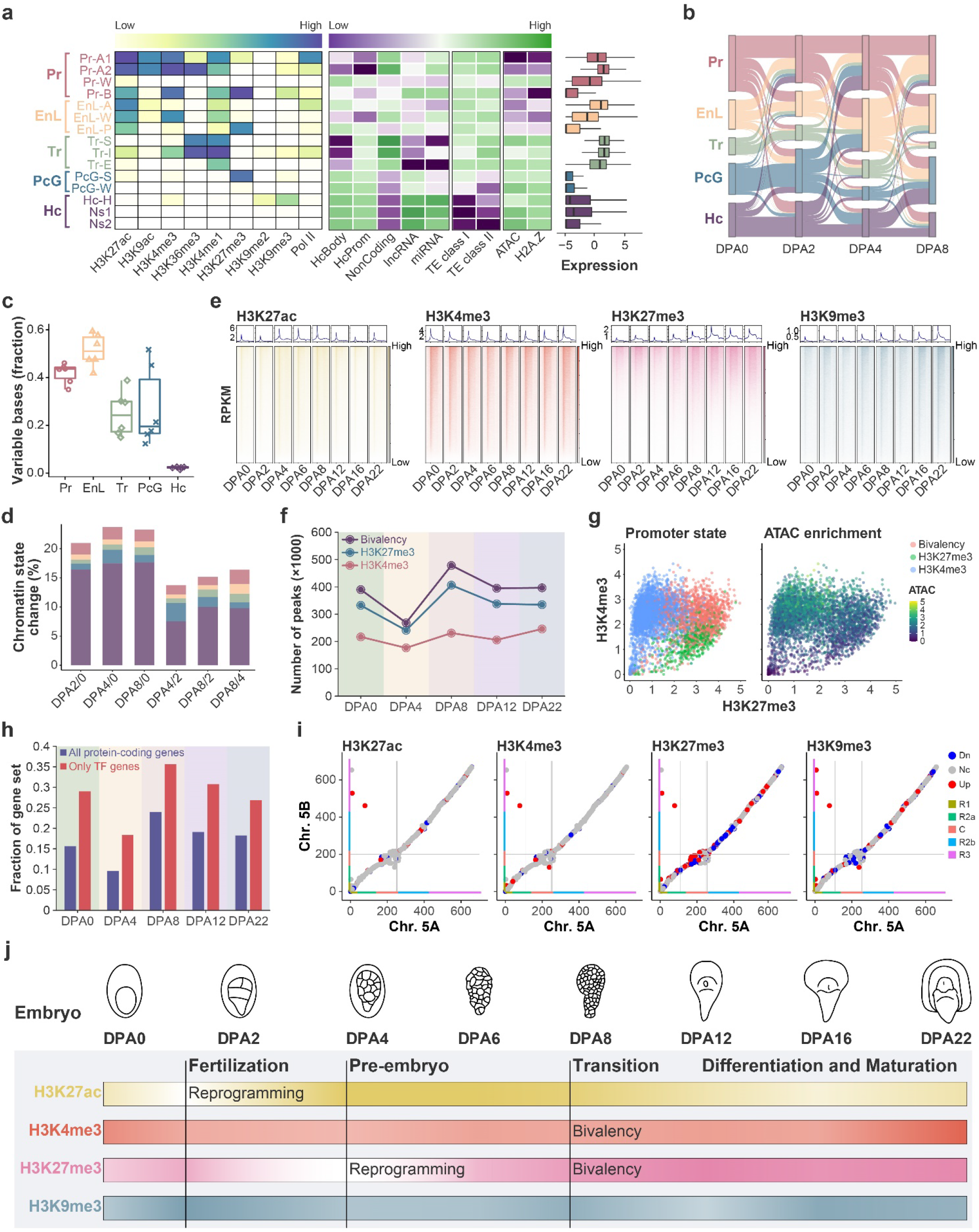
Reprogramming of histone modifications during wheat embryogenesis. **a**, Emission probabilities for histone modifications in 15 ChromHMM states, which could be mainly categorized in five groups: Pr (promoter), EnL (enhancer-like), Tr (transcription), PcG (polycomb group), and Hc (heterochromatin) state. The abbreviation following dash: A: active; W: weak; B: bivalency; P: poised; S: strong; I: initiation; E: elongation; H: H3K9me-associated; Ns: No signal. **b**, Chromatin states transition among different developmental stages. **c**, Dynamic of five major chromatin states during different embryonic stages. **d**, Percentage of changed chromatin state coverage to the total genome size among different embryonic stages. Colors filled represent the five major chromatin state groups as shown in **a**. **e,** Histone modification dynamic in promoter and genic regions during wheat embryogenesis. **f**, Regions of H3K4me3, H3K27me3 and bivalency at different developmental stages. **g,** Scatter plots correlating H3K27me3 (x axis) and H3K4me3 (y axis) enrichments (log_2_CPM) at proximal region. Different colors in the left panel indicate various modification states as labeled and chromatin accessibility of correlated plots is shown in the right panel with color bar for relative intensity (log_2_CPM). **h,** Fractions of all protein-coding and TF-coding genes marked by bivalent promoter at five major developmental stages. **i, Co**mparisons of histone modifications H3K27ac, H3K4me3, H3K27me3 and H3K9me3 of collinear regions between Chr. 5A and Chr. 5B. Down- (Dn), and Up-regulated (Up) loci were defined by the log_2_(fold change Chr. 5B/ Chr. 5A) values bigger and lower than 1 and -1, respectively. Nc indicates no change. Color bars and grey lines indicate the chromosome segments and centromere regions as those in Fig. 2d. **j,** Schematic represents the chromatin state dynamic during wheat embryogenesis. The dark and light colors represent the strength and weakness of the signal, respectively.

In mice and humans, histone modifications such as H3K4me3, H3K27me3, and H3K9me3 are extensively reprogrammed after fertilization and pivotal for early embryogenesis^4, 16^. Thus, we take a close look at their behaviors in wheat embryonic development. Expect for H3K36me3 and H3K4me1, most of histone modifications examined show a heavy proportion distributed in the distal intergenic regions (Supplementary Fig. 2a). We investigated the dynamic profiles of each histone modification with separation of proximal and distal regions. Different from animals, genome-wide H3K4me3 showed a slight change during embryogenesis of wheat at both proximal and distal regions (Fig. 3e and Supplementary Fig. 5a). Whereas, H3K27ac presented a quick decline after fertilization, then restore and maintained high till mid-embryo followed by gradually decreasing at late and mature embryo stages (Fig. 3e and Supplementary Fig. 5a). For H3K27me3, instead of being erased quickly after fertilization like the case in animals, a sharp erasure occurred at pre-embryo stage then gradually restored and kept high at late and maturation stages. H3K9me3 fluctuates slightly after fertilization and during pre-embryo, and then increased through mid-embryo till maturation (Fig. 3e and Supplementary Fig. 5a). Besides the individual histone modification, H3K4me3 and H3K27me3 co-occupied regions are referred as bivalent chromatin, which is correlated with and function in lineage-specific genes activation or silencing during embryonic development in animals^4, 16^. During wheat embryogenesis, bivalency reached its peak at transition stage (DPA 8) and gradually decreased during tissue differentiation (Fig. 3f). This is mostly due to the turbulence of H3K27me3, since H3K4me3 generally didn’t change much (Fig. 3f). In both proximal and distal bivalent regions, we observed widespread ATAC-seq, H3K27ac, H3K4me1, H3K9me3, Pol II and H2A.Z signals, but not transcript products and transcription-associated H3K36me3, suggesting a poised state (Fig. 3g). Of note, genes encoding transcription factors are more enriched than others in bivalent regions at every embryonic stage, indicating that H3K27me3 and bivalency are indispensable for accurate transcriptional regulatory circuitry (Fig. 3h). Besides, subgenomes of hexaploid wheat showed a different abundance of histone modifications even at collinear regions. Interestingly, H3K27me3 and H3K9me3 exhibited higher variation among different sub-genomes than active histone marks such as H3K27ac and H3K4me3, as well as histone variant H2A.Z (Fig. 3i and Supplementary Fig. 5b).

Taken together, the histone modification reprogramming is distinct in plants as compared to their counterparts in animals (Fig. 3j), indicating a divergent function of individual histone marks in driving the embryogenesis process in two kingdoms. As well, different subgenomes have varied histone modification, especially H3K27me3.

### Chromatin regulation of maternal gene silence and zygotic gene activation upon fertilization

Upon fertilization, differentiated parental gametes fused and converted into a totipotent zygote, including maternal gene silencing and zygotic genes initiation with epigenetic reconfiguration in both animals and higher plants^8, 15, 43^. A large amount of differential expressed genes (DEG) was captured upon fertilization (DPA2/ DPA0), with 8,079 down-regulated genes, mainly involved in maternal tissue identity maintaining such as AP2 and NF-YC9, as well as 8,380 up-regulated genes, including cell cycle and cytokine (CK) signaling genes associated with zygotic initiation (Fig. 4a). We next investigate the chromatin landscapes for seeking of driving force for those DEGs. Consistently, active histone modification H3K27ac and H3K4me3 suffered an overall lopsided decrease at down-regulated genes, while nearly no change at up-regulated genes (Fig. 4b). By contrast, chromatin accessibility didn’t change much for down-regulated genes but gained at up-regulated genes (Fig. 4b). As for H3K27me3, regardless of up or down-regulated genes, a reduction upon fertilization was observed. Such profiles indicated that loss of active histone modification might contribute to genes down-regulation, while gain of chromatin accessibility may trigger gene activation. Indeed, attenuation of either or both H3K27ac and H3K4me3 is extensively enriched for down-regulated genes (Fig. 4c, d), with more remarkable down-regulation for loss of both. H3K27ac reprogramming is more determinable compared with H3K4me3 (Fig. 4c, d). Genes subjected to both active histone marks decreasing and transcription levels attenuation are mainly maternal silenced genes, such as orthologs of floral homeotic gene AP2, signaling-related gene MAPKKK17, and stigma expressed gene IAA1 (Fig. 4e, h). Whereas, *de novo* building of pACRs showed a significant overlap and correlation with genes up regulated (Fig. 4f, g). These genes were enriched for pathways in DNA replication, cell cycle, heterochromatin, etc., which are the cornerstones of the early zygotic building (Fig. 4e, h).

**Fig. 4.**
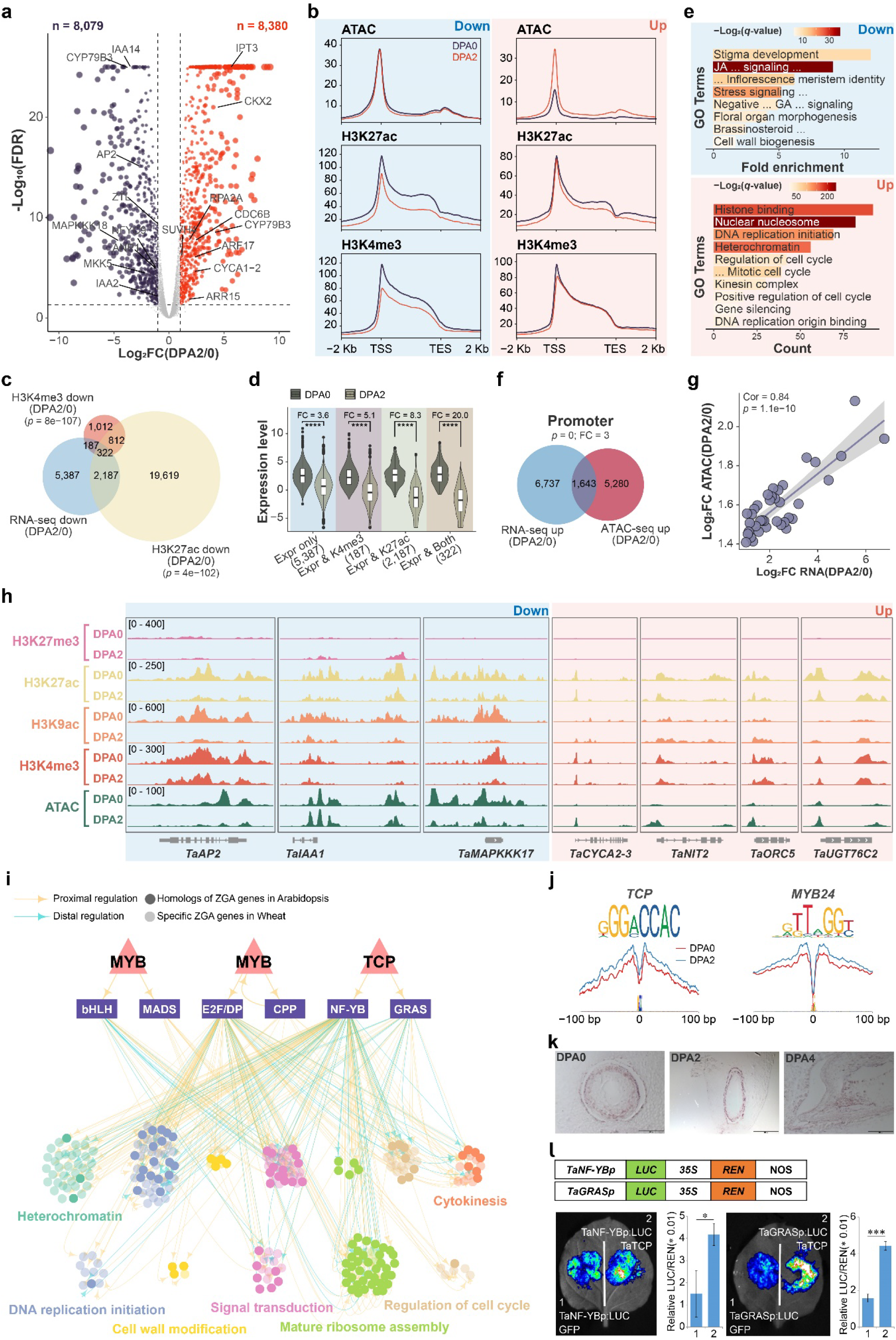
Transcription and chromatin reconfiguration during MZT. **a**, Volcano plot of differentially expressed genes during MZT. DEGs were defined by the threshold value of log_2_(Fold Change DPA2/DAP0) ≥1 and FDR ≤0.05 by edgeR. **b**, Chromatin accessibility and histone modification profiles of down- (left) and up-regulated (right) genes. **c**, Overlap between down-regulated genes and corresponding active histone modification marks decreasing. **d**, Expression levels of different gene sets generated from **c**. **e**, GO enrichment for 2,696 down-regulated genes in which active histone modification marks decreased (**c**) (top) and 1,643 up-regulated genes in which gained pACRs (**f**) (bottom). **f**, Overlap between up-regulated genes and corresponding chromatin accessibility gains. **g**, Correlation between gained pACRs and up-regulated genes based on 1643 genes overlapped. Genes were ranked by RNA-seq fold change and separated into 50 bins. **h**, IGV showing histone modifications and chromatin accessibility changes on representative down-regulated genes *AP2*, *MAPKKK17*, as well as *IAA1* (light blue shade), and up-regulated genes *CYCA2-3*, *NIT2*, *ORC5*, as well as *UGT76C2* (light red shade). **i**, GRNs for hierarchical regulation of zygotic active genes sets. Both proximal (orange) and distal (green) regulations were included. The Upper and lower layers of GRN represented the predicted regulation hierarchy. Each point represented a TF or gene. The target genes with orthologs of Arabidopsis ZGA genes were represented in light color, while the specific genes were represented in dark color. **j**, ATAC-seq footprint of key TFs in **i** during MZT. **k**, Expression patterns of *TCP* by *in situ* at DPA 0, 2 and 4. **l**, Luciferase reporter assays validation of transcriptional activation capability of TCP on targets *TaNF-YB* and *TaGRAS*. Hypergeometric distribution test was used for **c** and **f**. Mann-Whitney U test (two-sided) and Student’s t test were used for **d** and **l**, respectively (* : p <= 0.05; ** : p <= 0.01; ***: p <= 0.001; ****: p <= 0.0001). Pearson correlation test was used in **g**.

Besides, there are still 6,737 genes, nearly 80% of up-regulated genes upon fertilization, which can’t be explained by either chromatin accessibility or histone modification change (Fig. 4f). We thus speculate a hierarchy transcription regulation network (TRN) might govern the sequential activation of ZGA genes, like the case in animals^44, 45^. We constructed such TRN based on top-down and bottom-up methods via integrating the ATACseq and RNAseq data (Method; Fig. 4i and Supplementary Fig. 6b). Three TFs, one TCP and two MYBs were on the top layer of the pyramid (Fig. 4i). The footprint of TCP and MYBs showed not only an augmenting binding strength after fertilization but also transcriptional activators’ character, generating robust flanking accessibility without change of the footprint depth (Fig. 4j)^38^. In addition, the basal targets, which were activated at DPA2 or DPA4, were extremely conserved with more than 80% of which have orthologs in *Arabidopsis* ZGA genes set^8^, including genes involved in heterochromatin, cell wall, and cytokinesis (Fig. 4i). We further verified the specific expression pattern of TCP during MZT process by in situ hybridization (Fig. 4k), and its activation of downstream targets *GRAS/NF-YB* by reporter assay (Fig. 4l).

Taken together, the silence of the maternal genome is attributed to the diminishing of active modification on genes, while the trigger of zygotic may own to *de novo* building of chromatin accessibility and hierarchical regulation of TFs, with several potential key TFs such as TCP being engaged.

### Reprogramming of H3K27me3 coordinates with chromatin accessibility to ensure pre-embryo development

Following fertilization, several cell fate determination genes were gradually activated in pre-embryo, such as genes in WOX and ARR families (Fig. 1c, d). In parallel, we observed a dramatic drop of H3K27me3 at DPA4, but progressively re-build until DPA8 (Fig. 3e, f, j, Fig. 5a and Supplementary Fig. 7a). For developmental essential genes such as *LEC1*, *BBM*, *WOX11* and *ARR12*, local chromatin environments were permissive in pre-embryo (DPA4), with decreasing of repressive mark H3K27me3 and increasing of active marks H3K27ac and ATAC signal, while a limited change of H3K4me3 (Fig. 5b). What is more, about 90% of the H3K27me3 erasure regions at pre-embryo were re-built in transition stage, along with a large amount of *de novo* gain of H3K27me3 regions (Fig. 5c). The erasure of H3K27me3 was associated with CK pathway genes, embryo essential genes, and tissue specification genes, while the few augment was involved in auxin-related gene sets (Fig. 5c and Supplementary Fig. 7b, c). Correlated with the massive gain of H3K27me3 at DPA8, TaCLFs, a methyltransferase for H3K27, were temporally activated around DPA8 (Supplementary Fig. 7d), and several functional tested ‘PRE’ type motifs^46, 47^ were enriched at H3K27me3 restoration loci with their cognate binding TFs highly expressed at DPA8 (Supplementary Fig. 7e, f). Thus, cis-motif and trans-factors mediated H3K27me3 deposition may be involved in embryonic development in wheat.

**Fig. 5.**
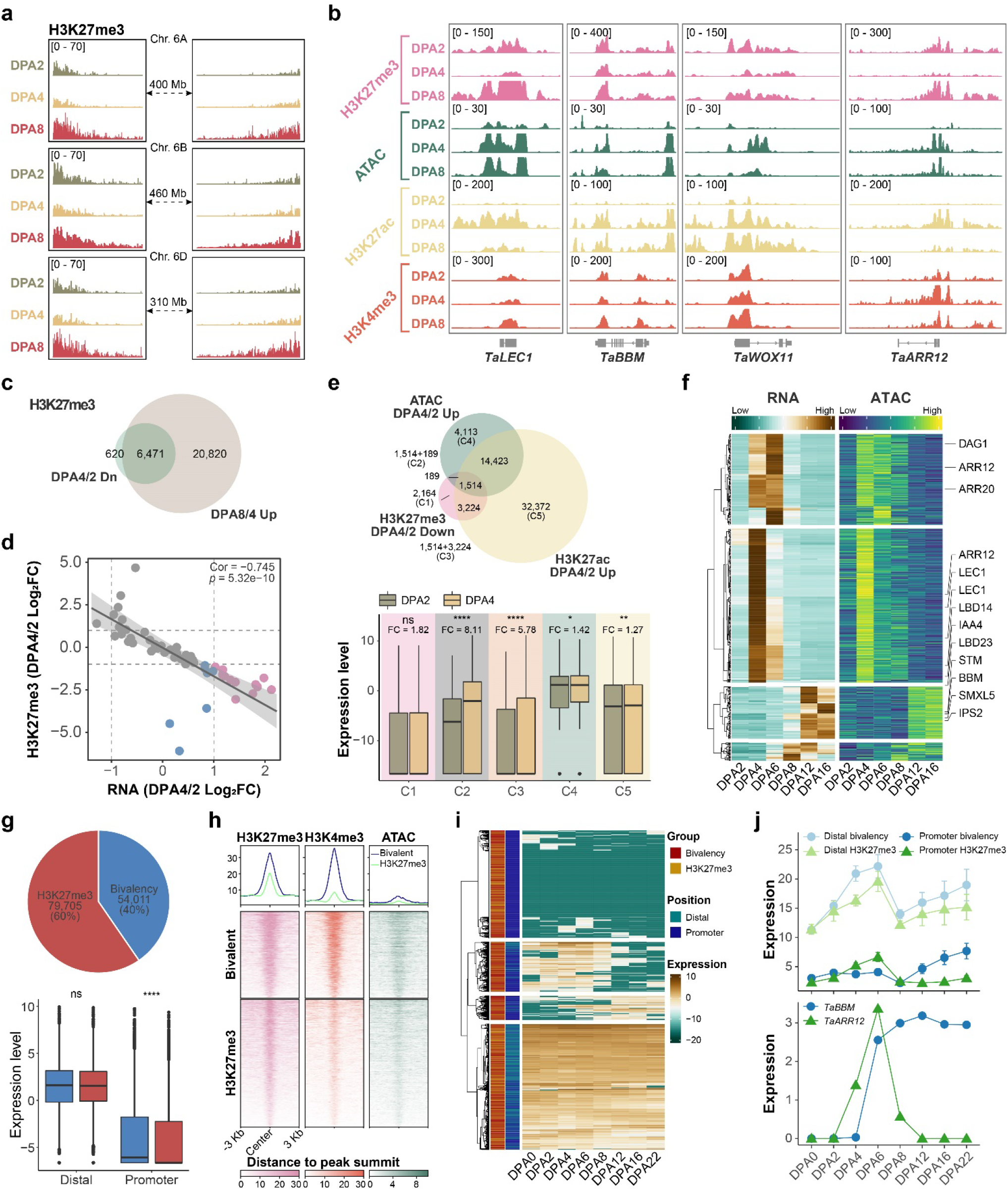
H3K27me3 reprogramming ensures pre-embryo development. **a**, Dynamic of H3K27me3 modification in Chr. 6 during embryogenesis, with resetting at DPA4 and regaining at DAP8. **b**, IGV showing histone modifications and chromatin accessibility changes on embryo developmental essential genes. **c**, Overlap between genes with H3K27me3 decreasing at DPA4 and increasing at DPA8. **d**, Correlation between H3K27me3 decreasing and target genes’ activation from DAP4 to DPA2. Genes were ranked by RNA-seq fold change and separated into 50 bins. H3K27me3 loss with mRNA level increased genes are shown in red color, while loss of H3K27me3 without mRNA changed is shown in blue color, while gray color indicates no change of H3K27me3. **e,** Overlap among genes with down-regulated H3K27me3 and up-regulated ATAC-seq and H3K27ac modification (top), and altered expression levels of different gene sets (bottom). **f,** Synchronous pattern between the gain of chromatin accessibility and elevation of gene expression on the basis of H3K27me3 decreasing at DPA4. Several developmental essential genes were highlighted. **g**, Proportion of H3K27me3 lonely and bivalent regions following H3K27me3 increasing at DPA8 (top), and the expression level of corresponding target genes. Distal targets assignment was based on reg-dACR-target links from Fig. 2h. **h**, Histone modifications and ATAC-seq enrichment at H3K27me3 lonely or bivalent regions. **i**.**j**, Expression pattern of genes associated with H3K27me3 lonely or bivalent modification across wheat embryogenesis. Heatmap showing clustering of gene expression (i), Line chart indicates the mean expression of various types of genes as indicated (top), as well as representative genes *TaBBM* (promoter bivalency) and *TaARR12* (promoter H3K27me3 alone) were shown (bottom) (**j**).

H3K27me3 intensity generally showed a negative correlation with transcription (Fig. 5d and Supplementary Fig. 2b), however, decreasing of H3K27me3 alone could not guarantee gene activation from DPA2 to DPA4 (Fig. 5e). However, H3K27me3 reduction was highly overlapped with the addition of H3K27ac or chromatin accessibility (Fig. 5e). As expected, genes with a gain of active chromatin modification in addition to H3K27me3 reduction showed significant activation, with a more dramatic effect by a gain of chromatin accessibility (Fig. 5e). Furthermore, there were 561 genes whose transcriptional activation showed highly synchronous patterns with the gain-of-chromatin after DPA4 (Fig. 5f and Supplementary Fig. 7g). This synchronization ensured the proper timing of genes activation, with embryo essential gene *LEC1* firstly activated followed by pluripotency gene *ARR12*, and later differentiation genes *XMXL5* and *IPS2* (Fig. 5f and Supplementary Fig. 7g). Together, this suggested that resetting of H3K27me3 lifted the chromatin layer barrier on embryo patterning genes, which was progressively activated along with the gain of chromatin accessibility. Alternatively, a global regain of H3K27me3 was observed at DPA8, a transition stage with the maximal open chromatin and bivalent regions as well (Fig. 1f, 3f, 5a, c, g and Supplementary Fig. 7a). Indeed, among the 133,716 gained H3K27me3 peaks, 40% show bivalency status with enriched H3K4me3 and opened chromatin status (Fig. 5g, h). The expression levels of genes with H3K27me3 marked promoters were significantly lower compared with bivalency promoters at transition stage, while slight change was detected between genes with distal H3K27me3 and bivalency (Fig. 5g, i). Genes got bivalency or H3K27me3 at both promoter and distal regions were silenced at transition stage, with stronger suppression function of modifications at promoter compared with those at distal (Fig. 5g, i, j). Yet, genes with bivalent promoters still have the opportunity to re-activation, such as *BBM*, but not genes with H3K27me3, such as *ARR12* (Fig. 5i, j).

Thus, the global depletion of H3K27me3 in pre-embryo resulted in a permissive chromatin environment, where the timely opening of chromatin can trigger the activation of developmental genes. Bivalency modification generated by H3K27me3 restoring marked cell fate determination genes, whose dysregulation was related to transition of totipotent pre-embryo to differentiation.

### Gene regulatory network orchestrates embryo pattern formation

After transition stage, embryo pattern formation was initiated, along with precisely programmed transcription regulation pyramid^48^. The complex relationship in TF-genes and TF *per se* could be well exemplified by the continuous transcription trajectories (Fig. 6a, b). To test how the architecture of embryo patterning from totipotent stem cell to maturation, we calculated the pseudotime of stages covering early- to mature-embryo based on the PCA distance (Fig. 6b). As expected, an excellent association between gene expression and both pACR and dACR was observed (Fig. 6c), which indicated that the expression pattern of these genes was governed by individual ACRs and TFs binding. Comparison of genes expressed in the early time point (Cluster 1 and 2) and the later ones (Cluster 3 and 4) presented a functional diversity, of which the early elevated genes were involved in cell division, while the later ones in cell specification (Fig. 6d). What is more, motifs of TFs binding sites, whose orthologs can facilitate inducement of cell totipotency in *Arabidopsis*, were enriched in early build ACRs, such as the binding motifs of LEC1, MYB118, and WUS, etc. Whereas the TFs, which function in seed dormancy, mainly occupied the later elevated ACRs, such as ABI5 (Fig. 6e). This result demonstrated evolutionary conservation of several key factors in embryo patterning in monocot and eudicot.

**Fig. 6.**
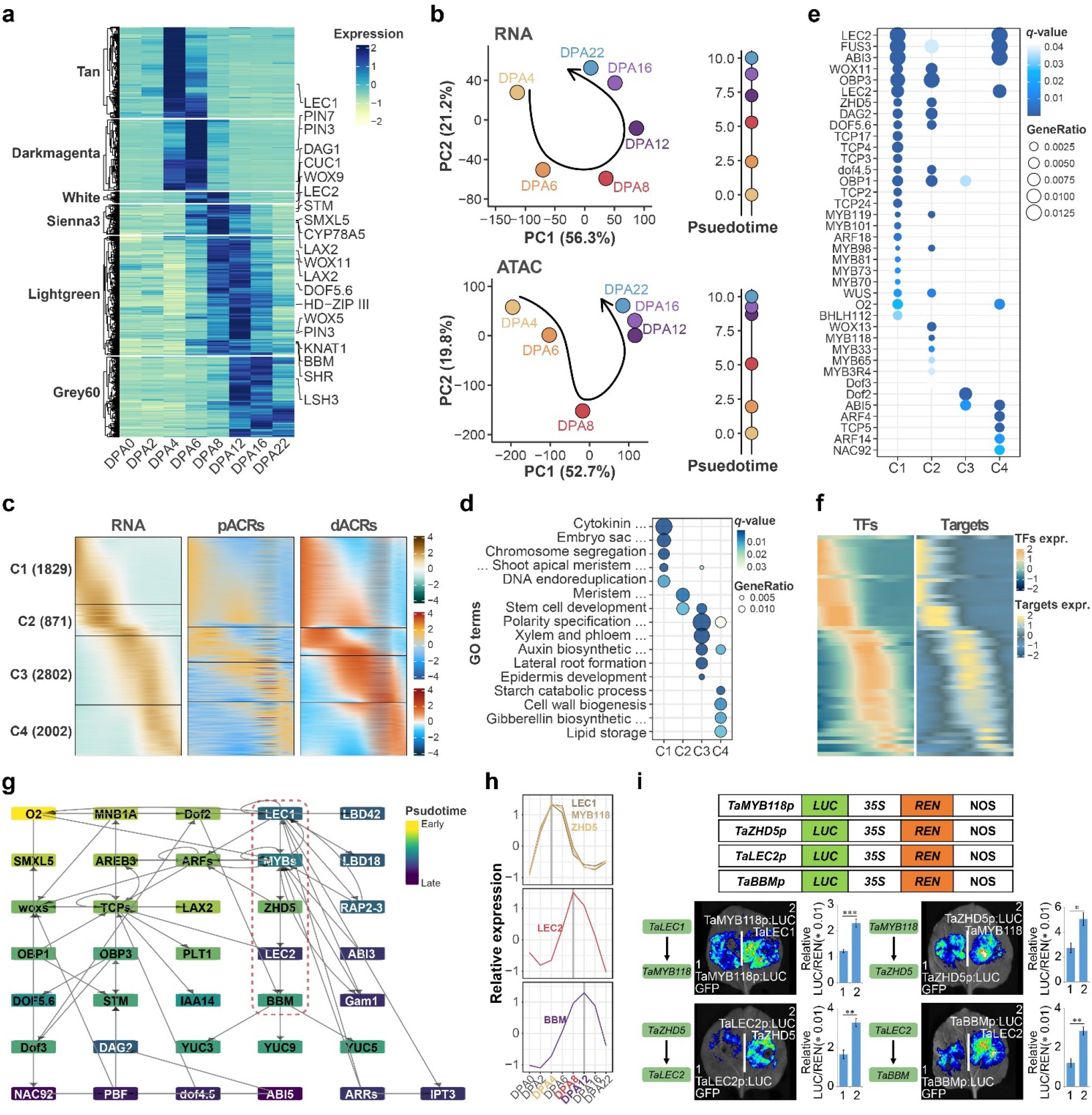
Transcription regulatory network governing embryonic patterning. **a**, Expression heatmap of selected key modules genes from Fig.1c. Genes with ortholog identified to function during embryogenesis in other species were indicated. **b**, PCA trajectories of genes expression (top) from (a) and corresponding chromatin accessibility (bottom). Developmental time units (DTU) values on the right panel were calculated based on the straight distance between two adjacent points and then scaled to 0 to 10. **c**, Sorted standardized temporal genes, pACRs, dACRs and expression profiles. Four clusters were generated based on expression order. **d,e**, GO and TFs enrichment for four clusters of genes from (**c**). **f,** Heatmap showing the synchronous patterns between TFs and target genes. Each row in heatmap represents links between one TF and corresponding genes. **g**, GRNs for key TFs participated in embryonic patterning. Genes were roughly ranked by the expression order from top to bottom and from left to right. Colors indicate the expression timing wave during embryo patterning. **h**, Expression pattern of representative key genes in the GRN network. **i**, Luciferase reporter assays validation of transcriptional regulation among representative TF-target pairs. Student’s t test was used for the statistic significance (* : p <= 0.05; ** : p <= 0.01; ***: p <= 0.001).

We further dissect the architecture of the gene regulation network (GRN). In addition to binding motif specificity, we filtered target genes to TFs by co-expression pattern as well (Method; Fig. 6f). As a result, 1,158 paired TF-cis motif interactions were identified including 154 target genes, 695 ACRs and 191 TFs (Fig. 6f). To better understand how TFs are coordinated during embryo pattern formation, we next extract the TF-TF regulation network (Fig. 6g). We found four pivotal classes of TFs located at the center of the network, TCPs, ARFs, MYBs, and WOXs. Some of those TFs such as WOXs and ARFs are reported to regulate embryo patterning in other species^49–51^. Several known genes regulating organ and tissue specification in Arabidopsis were under the governing of these TF modules, such as I*AA14, LAX2, PLT1,* and *SMXL5*, etc. To further estimate this GRN, we use in situ hybridization to verify the spatiotemporal expression pattern of key TFs module including LEC1-Myb118-ZHD5-LEC2-BBM. As shown, the *LEC1, Myb118* and *ZHD15* genes were highly expressed in pre-embryo stages of overlapped tissue, while *LEC2* was activated later, with a peak at transition stage, and *BBM* gradually induced at mid-embryo, indicating a sequential pattern (Fig. 6h). The luciferase activity from the reporter assay validated the regulatory circuit along LEC1, MYB118, ZHD5, LEC2 and BBM module (Fig. 6i). Interestingly, TaLEC2 could bind the pACR of *BBM* directly and activate it *in vitro* (Fig. 6i), which is not reported in the model plant *Arabidopsis*.

Thus, these data described a cohesive, sequential wave of TFs regulation in governing wheat embryo pattern formation.

### Chromatin regulation facilitates embryo maturation and prohibits extensive organogenesis

During seed maturation, nutrient was accumulated in both endosperm and embryo, meanwhile, the dormancy program was progressively established with increased ABA levels accompanied by desiccation^52^. Indeed, genes elevated specifically in late embryo development were enriched in carbohydrate, lipid, protein, amino acid, secondary metabolic pathway, and ABA signaling and homeostasis (Fig. 7a). As expected, H3K27ac increased progressively at those genes during late embryo development, but H3K27me3 declined to a relatively low level, while chromatin accessibility was slightly changed (Fig. 7b). Next, we applied ATAC footprint to seek candidate TFs associated with the regulatory regions of the above genes. ABA-related TFs were enriched for regulating ABA pathway genes, which verified the logic of this analysis (Fig. 7c). In addition, most nutrient storage pathways shared at least one upstream TF with ABA pathway, such as ABI5 and ABF1 (Fig. 7d), which are known to regulate a subset of late embryogenesis abundant genes in *Arabidopsis*^53^. This result suggested that several key regulators could synchronize nutrient accumulation and embryo dormancy in cooperation with chromatin landscape dynamics.

**Fig. 7.**
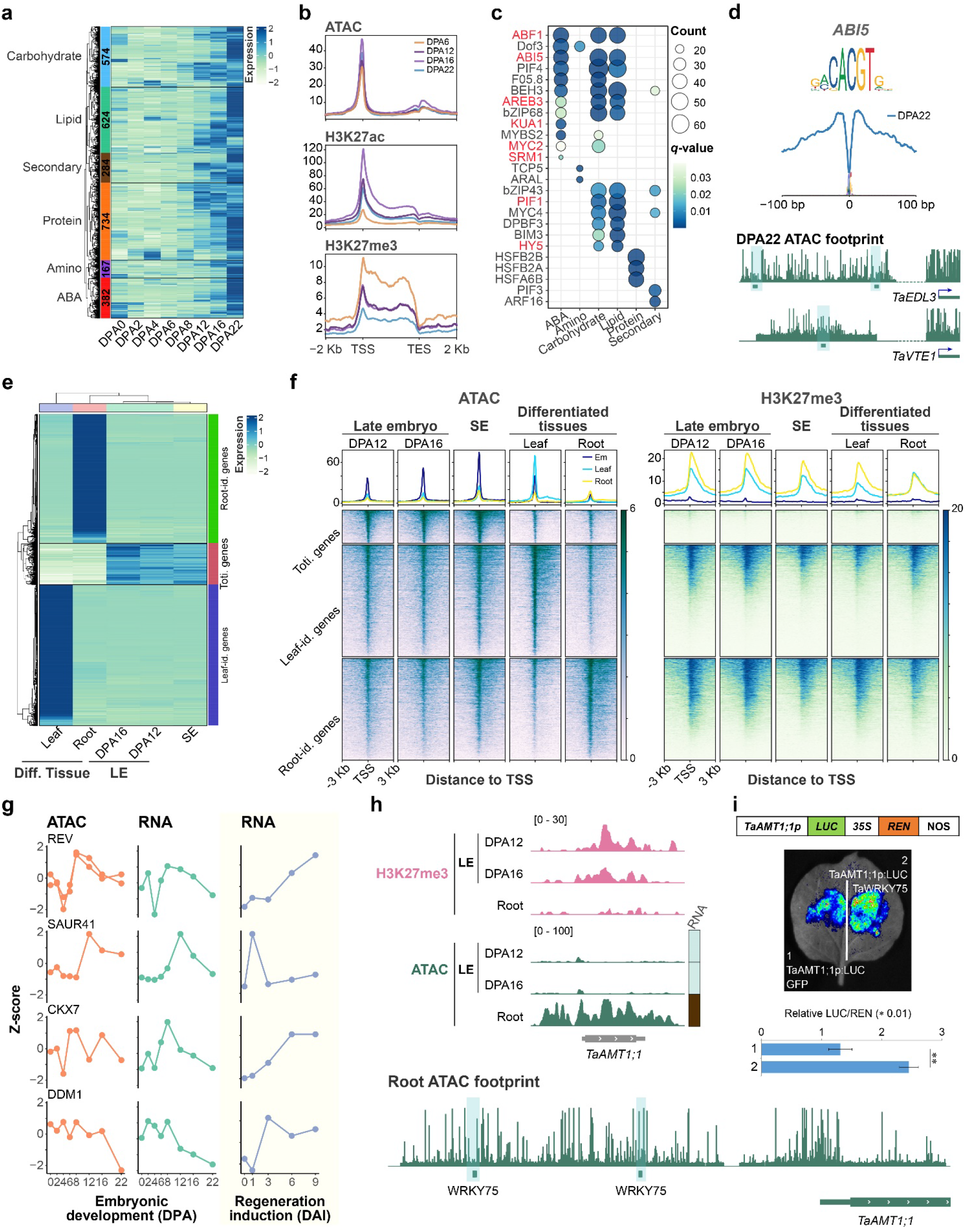
Maturation of embryogenesis in the context of chromatin regulation. **a**, **b**, Six pathway genes highly expressed during the late embryonic development (**a**), and corresponding epigenetic modifications (**b**). **c**, **d**, Regulation of six pathway genes by specific cis motif and trans-factors. Binding TF enriched for each pathway genes (**c**). ABI5 footprint was highlighted (**d** top), and corresponding binding site at the promoters of carbohydrate-related gene *TaEDL3* and ABA-related gene *TaVTE1* (**d** bottom). **e**, Identification of organ identity and totipotent genes based on expression specification across different tissues. **f,** ATAC and H3K27me3 modification on the promoters of different genes sets in **e** cross five tissues, SE: somatic embryo. **g**, Chromatin accessibility (left) and expression (mid) of regeneration genes during wheat embryogenesis, and their expression during callus induction (right). **h**, H3K27me3 and chromatin accessibility regulated root-identity gene *TaAMT1;1* specific expression pattern in root and late embryo (LE). The ATAC footprint tracks showing the binding sites of TaWRKY75, a key TF predicted to bind *TaAMT1;1* as indicated in supplementary Fig. 8e. **i**, Luciferase reporter assays show the transcriptional activation capability of TaWRKY75 to *TaAMT1;1* in *N. benthamiana* leaves. Student’s t test was used for the statistic significance (* : p <= 0.05; ** : p <= 0.01; ***: p <= 0.001).

Unlike animals, extensive organogenesis is prohibited during late embryo development, which enables plants to generate post-embryonic lateral organs in adaptation to changeable environments. Meanwhile, immature embryo served as excellent explants for wheat regeneration, a process including regaining of totipotency and organogenesis^53, 54^, however such capability was reduced upon embryo maturation (Supplementary Fig. 8a). We wonder how the totipotency capability is attenuated and why the terminal organ is not formed during embryo maturation. First, we identified organ identity genes by their specific expression in terminal organs such as root and leaf, for comparison totipotent genes were characterized with high expression in mid-embryo or embryonic callus (Fig. 7e). GO enrichment of different groups of genes verified the accuracy of genes identity (Supplementary Fig. 8b). Next, we evaluated the chromatin landscape of each cluster genes in different organs. Depletion of H3K27me3 on organ identity genes was observed at the respective organ, whereas, an obvious ATAC signal presented for both totipotency and organ identity genes in its corresponding organ (Fig. 7f, Supplementary Fig. 8c, d). Surprisingly, low/no H3K27me3 was enriched on totipotent genes even in differentiated organs (Fig. 7f). In embryonic callus, of which new plant will regenerate, the promoter region of both totipotent genes and organ identity genes were opened with high chromatin accessibility and low H3K27me3 level (Fig. 7f). Thus, chromatin accessibility and H3K27me3 coordinately regulated organ identity gene’s activation potential while chromatin accessibility but not H3K27me3 mediated repression is indispensable for totipotency gene silencing. Indeed, several regeneration-related factors, such as REV, SAUR41, CKX7 and DDM1 were gradually diminished during embryo maturation along with closing chromatin status but activated during callus induction process (Fig. 7g). Last, we investigated the gene-regulatory logic underlying plant organ formation. WRKYs and AtHB TFs footprint were enriched in regulatory regions of root and leaf identity genes respectively, whereas, bHLH and GATA TFs footprint were enriched in both gene-sets (Supplementary Fig. 8e). Remarkably, nearly half of the identified TFs did not show organ-specific expression, but instead a wide-spectrum pattern, indicating that TFs expression *per se* cannot determine the specificity of organ identity (Supplementary Fig. 8f). For example, although WRKY75 was expressed during embryo maturation and it owns the capability to bind and active *AMT1;1,* an ammonium transporter coding gene in root (Fig. 7h, Supplementary Fig. 8f), AMT1;1 was silenced due to the H3K27me3 repression and compaction chromatin environment at the promoter (Fig. 7i).

Taken together, chromatin layer regulation ensures the proper embryo maturation process, with an accumulation of nutrients and reduced totipotency ability but not extensive organogenesis.

### Epigenetic regulated ontogenetic divergence and inverse hourglass pattern during embryogenesis

As wheat is an allohexaploid plant, we further extend the understanding of epigenetic regulated embryogenesis to the evolution and polyploidization perspectives. Genes were clustered based on their evolutionary age (Supplementary Fig. 10a). Most of the genus *Triticum*-specific genes, especially D-subgenome unique genes, were located within PcG chromatin state and did not participate in embryonic development (Supplementary Fig. 10a-c). For the homeolog triads that are present in all three subgenomes, chromatin accessibility is related to the bias expression pattern (Supplementary Fig. 11) in addition to histone modifications as reported^55^. In detail, suppressed triads are generally more than dominant triads, in addition, D-subgenome suppressed triads are relatively less than A or B-subgenome suppressed. More balanced triads were expressed in the middle period compared with early and late embryogenesis (Supplementary Fig. 10d). Such observation was generally consistent with the previous report but different in details^27^.

Polyploidy may drive genes neo- or sub-function to confer phenotypic plasticity, which can be represented by the varied expression pattern. Instead of identifying genes of differential expression at a specific embryonic stage by comparing hexaploid wheat with tetraploid and diploid^27^, we look for genes showing varied expression profiles during the process of embryo development. Indexing by Pearson correlation degree, genes are categorized into dysfunction, middle, and conserved clusters (Fig 8a and Supplementary 9a). As expected, conserved genes were mostly homeologs triads, with sub-genome balanced expression, and showed higher sequence conservation and stronger selection pressure (Fig. 8b, c and Supplementary Fig. 9b-f). However, a considerable proportion of dysfunctional genes changed expression profile in hexaploid wheat as compared to ancestors, but with slight sequence variation and subject to strong negative selection (Fig. 8c). Promoters of these genes, especially around 3Kb upstream of TSS, were preferably inserted by TEs (Fig. 8d and Supplementary Fig. 9g). Intriguingly, TE insertion regions tended to be accessible chromatin and enriched for H3K27ac and H2A.Z (Fig. 8e). For example, TaUGT91C1 showed a B-subgenome dominance expression in hexaploid wheat, while silenced in all ancestors (Fig. 8f). Accordingly, the promoter and dACR region harbored TEs with higher chromatin accessibility as compared to A or D counterparts. Therefore, the TEs insertion within proximal and distal active chromatin regions may drive the novel expression pattern of offspring genes following polyploidization.

**Fig. 8.**
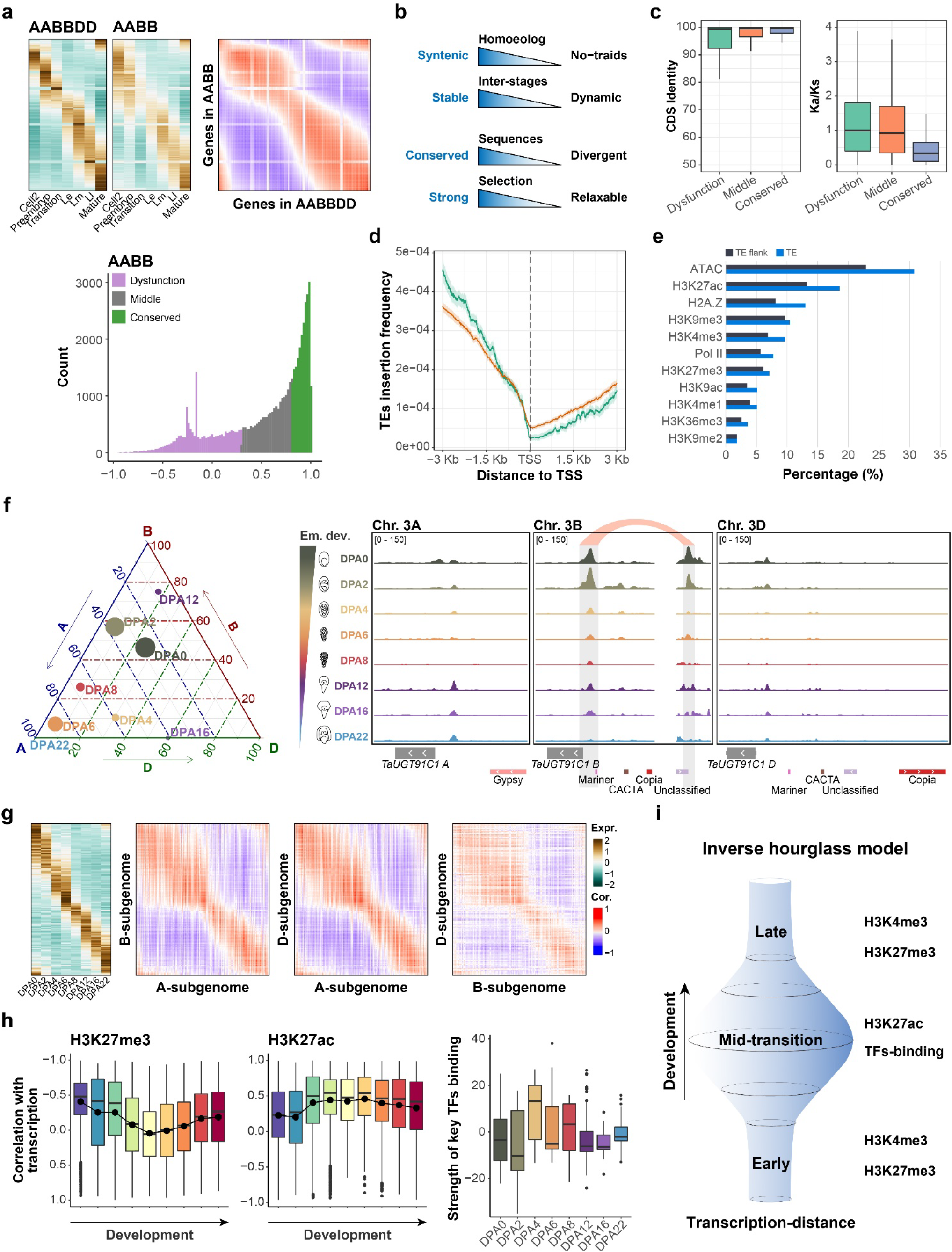
Transcriptional divergence between different ploidy wheat and epigenetic governed inverse hourglass model. **a**, Comparisons of gene expressions between hexaploid wheat (AABBDD) and ancestor (AABB). Each row represents an individual gene in hexaploid wheat and its counterpart ancestor (top left). Pearson correlation index was calculated between genes expression in hexaploid wheat and counterpart in ancestor (top right). Genes were clustered into three categories based on Pearson index (bottom). **b**, Genomic features differences between dysfunction and conserved cluster genes generated in **a**. Statistics were generated from supplementary Fig. 9b-f. **c**, Coding sequence similarity and Ka/Ks comparisons among different gene sets in **a**. **d**, TEs insertion frequency in the promoters of different gene sets in **a**. **e**, epigenetic modification enrichment at TEs insertion loci. **f,** Homoeologs bias expression of representative gene TaUGT91C1 (left) and epigenetic regulations (right). **g**, Comparisons of homoeologs expressions among subgenomes of hexaploid wheat. Each row represents an individual gene A-subgenome of hexaploid wheat, and the genes were ranked by activated order (left). Pearson correlation index was calculated between homoeologs expression in different subgenomes of hexaploid wheat as those in **a**. **h**, H3K27me3 and H3K27ac contribution to genes expression across wheat embryogenesis (left) and key TFs binding strength at different developmental stages (right). The top 100 variable motifs calculated by chrVAR were used as key TFs. **i,** The*‘*inverse hourglass model’. The transcription distance was large in the mid-stage. Histone modification H3K4me3 and H3K27me3 contributed to the early- and late-stage, and H3K27ac and TFs-binding contributed to the mid-stage.

A previous study on embryogenesis revealed a conserved hourglass model in hexaploid wheat as well as its tetraploid, and diploid ancestry, but a slight difference for the phylotypic stage^27^. Here, our transcriptome data in hexaploid wheat (Chinese Spring) also supported that transition stage of DPA8 represents the phylotypic stage (Supplementary Fig. 9h). In addition, we find an inverse hourglass model for transcriptional variability. A sharp mid-developmental transition was presented, which separated two conserved phases, early and late developmental stages in both the comparison between homoeologs in hexaploid *per se* and between counterparts of hexaploid gene and its ancestors (Fig. 8a, g and Supplementary Fig. 9a). We hypothesized that epigenetic regulation may account for this middle-stage transition. Indeed, H3K27ac was mainly contributed to the middle stage rather than early- and late-stage, whereas H3K27me3, H3K4me3, and ATAC behaved oppositely (Fig. 8h and F Supplementary Fig. 9i). In addition, chromVAR analysis indicated that the key TFs during embryogenesis functioned mainly in the middle stage (Fig. 8h and F Supplementary Fig. 9j). Thus, the cis- and trans-regulation driven by TFs was more meaningful in mid-stage compared with early- and late-stage. In sum, transcriptional variance showed an inverse hourglass model, which may be governed by chromatin layer regulation at early and late stage, while cis-trans regulation at the middle stage (Fig. 8i).

## DISCUSSION

Despite containing the same genome, different cell types possess unique identities with specific gene expression patterns. This is partly because some unique portions of the genome are accessible in each cell type, which is controlled at the level of chromatin. During embryogenesis, parental differentiated gametes fuse and convert into a totipotent zygote, and then to a differentiated embryo, cells undergo extensive identity switch, and thus provide an ideal system for studying chromatin-based transcription regulation. Indeed, embryogenesis is extensively studied in mammalians. However, our understanding of embryogenesis is far lagged in plants, especially in crops. In this study, we generated comprehensive RNAseq, CUT&Tag and ATACseq data in three early embryo sac and five later embryo samples of hexaploid wheat Chinese Spring. It provides a genome-wide annotation of dynamic ‘reference epigenome’, which enables the comparative analysis of large-scale chromatin reprogramming and its function for embryogenesis between plants and animals, and the understanding of multi-layer orchestrated transcriptional regulation circuit at different stages of embryonic development in wheat. Integration with the previous report on the transcriptome of embryogenesis from different ploidy wheat^27^, our study further revealed the potential driving force for evolutionary divergence among different sub-genome in shaping embryogenesis in polyploidy wheat.

In this study, we observed large-scale chromatin modification reprogramming during wheat embryogenesis, especially at MZT, pre-embryo and transition stages (Fig. 3e, j, Fig. 4b, Fig. 5a and Supplementary Fig. 5a). Though the histone modification resetting pattern shows general similarity to animals^4, 16^, there was a large variation in terms of modification types and timing of reprogramming. In wheat, widespread H3K27ac dropped off during MZT and contributed to the silence of maternal genes, slightly for H3K4me3 (Fig. 3e, j and Fig. 4b), which however is dramatically reset upon fertilization and re-build during major ZGA and is importan for ZGA in mice^4, 15^. H3K27me3 sharply dropped at pre-embryo but not right after fertilization like its counterpart in mammals^4^ (Fig. 3e, j, Fig. 5a and Supplementary Fig. 5a). Both H3K27ac and H3K27me3 were momentarily and universally re-established, of note, regions of H3K27me3 decreased in DPA4 is a great part of that increased in DPA8 (Fig. 3j, Fig. 5a and Supplementary Fig. 5a). This suggests the epigenome ‘rebooting’ model is conserved for both animals and plants^15^. Importantly, H3K27me3 dynamics marked embryonic essential genes (Fig. 3h and Fig. 5b), indicating its pivotal role in embryo development. Indeed, we found aberrant seed development when knocking out H3K27me3 writer complex component (Supplementary Fig. 12), though a detailed mechanism still needs to address. Recent findings characterize the direct involvement of PRC2 component, writer of H3K27me3, for embryo development in *Arabidopsis*^56^. Besides histone modification, chromatin accessibility also shows dynamics during embryogenesis. Interestingly, dACRs (account for 75% total ACRs) and pACRs (16%) show unique fluctuation pattern, correlated with distinct transcriptional regulation respectively (Fig. 2b). For genic regions, pACRs were significantly gained upon fertilization, strongly correlated with zygotic gene activation (Fig. 4b, e-h), and continued gain till pre-embryo (DPA4) then gradually declined later on. Whereas, dACRs gradually gained after fertilization, with ‘transient’ maximize at transition stage (DPA8) then quickly declined later, where the opened intergenic regions are highly overlapped with multiple TEs, such as Gypsy and CACTA, as well as ‘burst’ of H3K9me3 region at centromere and telomere. Further analysis of global DNA methylome and enhancer RNA profiling around transition stage would give a clearer conclusion of the functional consequence of such dACR eruptions. Nevertheless, our data provide a novel comparison between animals and plants for the whole embryogenesis process driven by chromatin landscape dynamics.

Embryogenesis is governed by precise transcriptional regulation program^2, 3, 57^. However, how such transcription regulation is orchestrated by different types of regulation, such as histone modification, chromatin accessibility, DNA methylation, transcription factor-cis motif modules, is not clear. Based on our analysis, a dynamic ‘three-layer regulation model’ is proposed for different embryonic development stages (Fig. 8i). In this model, de-repression of local histone modification is a prerequisite of permissive chromatin environment and open chromatin is the cornerstone of transcription factors free binding. At pre-embryo, H3K27me3 was globally reset, which provides a flexible chromatin environment, but yet, not sufficient to active genes and gain-of accessibility at the promoters are the indispensable triggers. After this, freestanding TFs coordinate was gradually built in a subtle and specific manner (Fig. 5). The attenuation of totipotency and prohibition of extensive organogenesis at late-embryo development stage can also support this model in an opposite way (Fig. 7e-i). A representative case is the WKRY75-*AMT1;1*. Despite the transcriptional activation of WKRY75 for *AMT1;1* in reporter assay, *AMT1;1* was still repressed in late-embryo because of the compacted promoter region along with enriched H3K27me3. Besides the stage-specific cases, similar multi-layer regulation could apply to the entire embryogenesis process in general. Transcription data for embryonic development demonstrate a cross-kingdom and ploidy conserved ‘inverse hourglass model’, in which a transcriptional divergent mid-developmental transition punctuated the conserved early- and late-stage (Fig. 8a, g and Supplementary Fig. 9a). Accordingly, a panorama of genetic and epigenetic dynamic presents that global histone modification reprogramming at early-stage, while the chromatin accessibility burst and TF-binding at mid-stage as well as the relative compacted genome at late-stage (Fig. 4-7, Fig. 8h-i and Supplementary Fig. 9i, j). Thus, the early- and late-stage were dominated by chromatin level restriction, while mid-stage was governed by TF-motif regulatory circuits along with open chromatin, of which the chromatin level regulation effect is minimized. This regulation pattern fits the transcriptional divergent ‘inverse hourglass’ in general, in combination with the genetic diversity ‘hourglass model’ for mediating conserved and diversified embryogenesis in different species.

Bread wheat (AABBDD) undergoes polyploidization and domestication, along with evolutionary divergence among different sub-genomes. Indeed, we observed dynamic biased expression of homeolog triads during embryogenesis, correlated with histone modification variations as well as chromatin accessibility differences among A,B, D sub-genome (Fig. 2d, Fig. 3i, Supplementary Fig 3d and Supplementary Fig 5b). In addition, a considerable proportion of genes showed varied expression profiles during embryogenesis in allohexaploid bread wheat compared with the diploid ancestors^27^ (Fig. 8a and Supplementary Fig 9a). Part of them possesses DNA variations within the genic regions, which may alter gene function and indirectly change their expression profile as an adaptation for the ‘new’ roles. Whereas other genes with slight or nearly no sequence variation at genic region still show altered expression patterns. Of note, promoters and distal regulatory regions of those genes contain asymmetric TE insertions among different sub-genomes, with a preference for sites of open chromatin region and enriched for active histone marks such as H3K27ac and H2A.Z. Thus, epigenetic landscape variation among different sub-genome, especially TE inserted active chromatin regions mediated transcriptional alteration, may drive the evolutionary divergence in shaping embryogenesis in polyploidy wheat.

Besides, our data also elucidated detailed transcriptional regulation circuits during ZGA and embryo pattern formation, as well as the maturation of embryo (Fig. 4, Fig. 6, Fig. 7). The hierarchy transcription regulation pyramids and underlying epigenetic regulation of key transcription factors during ZGA would be worth to test for the automatic embryo development program without fertilization. As well, the knowledge about programmed embryo pattern formation and gradually attenuation of totipotency would be valuable for studying wheat regeneration, during which immature embryo serves as the excellent explants. As a comprehensive study of embryonic chromatin landscape dynamics and transcriptome, our research is anticipated to facilitate gene functional study of wheat embryogenesis and plant breeding application, such as parthenogenesis and somatic embryogenesis.

## Methods

### Plant materials and growth conditions

A spring wheat (Triticum aestivum) Chines spring was used in this study. All plant was grown in a growth room under a 16h light/8h dark cycle with light intensity 1,000 μmol m−2 s−1, and temperature 15-26°C depending on growth stage. The stamens are removed before the pollen matures. Then we conduct artificial pollination and record the number of days to ensure the accurate time of seed development. Embryo and embryo sac at the specific developmental stage were sampled using RNA and nuclear extraction.

### RNA extraction

Total RNA was extracted using HiPure Plant RNA Mini Kit according to the manufacturer’s instructions (Magen, R4111-02), and libraries were sequenced using an Illumina Novaseq platform.

### CUT&Tag experiment

The CUT&Tag experiment was performed as reported with minor modifications^35^. For each replicate, the fresh samples were soaked in the HBM buffer (25 mM Tris-Cl pH 7.6, 0.44 M sucrose, 10 mM MgCl2, 0.1% Triton-X, 10 mM Beta-mercaptoethanol, 2 mM spermine, 1 mM PMSF, EDTA-free protease inhibitor cocktail), after chopped with a razor blade, the mixture was filtered through a 40-μm cell strainer. The crude nuclei was washed twice by HBB buffer (25 mM Tris-Cl pH 7.6, 0.44 M sucrose, 10 mM MgCl2, 0.1% Triton-X, 10 mM Beta-mercaptoethanol). Then the crude nuclei were stained with 4,6-diamidino-2-phenylindole and counted by hemocytometer.

Extracted nuclei (10,000) was resuspend softly in 50µl antibody buffer (20 mM HEPES pH 7.5; 150 mM NaCl; 0.5 mM Spermidine; 1× Protease inhibitor cocktail; 2 mM EDTA; 0.1% BSA) containing a 1:50 dilution of corresponding antibody. After overnight incubation in 4 °C, the primary antibody was removed by centrifuged (1,200g,3min), the nuclei were incubated in 50µl wash buffer (20 mM HEPES pH 7.5; 150 mM NaCl; 0.5 mM Spermidine; 1× Protease inhibitor cocktail) with secondary antibody (1:100; Guinea Pig anti-Rabbit IgG antibody) at 4°C for around 1-2 hour and then wash twice with wash buffer. A 1:100 dilution of pA-Tn5 complex was prepared in CT-300 buffer (20 mM HEPES pH 7.5; 300 mM NaCl; 0.5 mM Spermidine; 1× Protease inhibitor cocktail), after nuclei centrifuged, add 100µl mix and incubate 2-3h in 4°C. After incubation with pA-Tn5, wash twice with CT-300 buffer. Then incubate nuclei in 300 µl Tagmentation buffer (20 mM HEPES pH 7.5; 300 mM NaCl; 0.5 mM Spermidine; 1× Protease inhibitor cocktail; 10 mM MgCl2) in 37°C for 1h. To stop tagmentation reaction, add 10 µl 0.5M EDTA, 3 µl 10% SDS and 2.5 µl 20mg/ml Protease K, incubate 1h in 50 °C. The DNA was extracted with phenol:chloroform:isoamyl alcohol, precipitated with ethanol and resuspended in ddH2O. The library was amplified 17 cycles by Q5 high fidelity polymerase (NEB, M0491L), and purified by AMPure XP beads (Beckman, A63881). Finally, the library was sequenced using an Illumina Novaseq platform.

### ATAC-seq experiment

The method of nuclei extraction was performed as before. After checking the nuclear integrity, the nuclei extracted (50,000 per reaction) were incubated with the Tn5 transposase and tagmentation buffer at 37 °C for 30 min (Vazyme Biotech,TD501-01)^34^. After tagmentation, the DNA is purified by PCR purification kit (QIAGEN, 28106). PCR was performed to amplify the library for 9-12 cycles using the following PCR conditions: 72 °C for 5 min; 98 °C for 30 s; and thermocycling at 98 °C for 15 s, 63 °C for 30 s and 72 °C for 40 s; following by 72 °C 5 s. After the PCR reaction, libraries were purified with AMPure beads (Beckman, A63881).

### Bioinformatics Data Preprocessing and Alignment

All fastq data, including DNA and RNA sequencing, were generated based on Illumina Hiseq-PE150. Raw data were filtered by fastp (v0.20.0) with “--detect_adapter_for_pe” parameter for reads filter, low-quality bases trimming, and adapters removing^58^. Furthermore, the clean data was evaluated by fastqc software (v0.11.8) (https://github.com/s-andrews/FastQC) to ensure the high quality of reads.

Both DNA sequencing including resequencing, CUT&Tag, and ATAC-seq and RNA sequencing data were aligned based on *Triticum aestivum* (Chinese Spring) genome assembly (IWGSC RefSeq v1.0)^31^, which was downloaded from https://urgi.versailles.inra.fr/download/iwgsc/IWGSC_RefSeq_Assemblies/v1.0/. The IWGSC Annotation v1.1 was used as the gene annotation reference. For DNA sequencing data, BWA-MEM (v0.7.17) algorithm was used for alignment with “-M” parameter to avoid shorter split hits^59^. For RNA sequencing data, hisat2 (2.1.0) was applied for reads mapping with default parameters^60^.

### RNA-seq Data Processing and Expression Clustering

Sam files generated from hisat2 were converted to bam files without deduplication. FeatureCount v1.6.4 was used for reads quantity per gene^61^. An R package edgeR was used for DEGs (differentially expressed genes) examination, with a threshold absolute value of Log2 Fold Change ≥ 1 and FDR ≤ 0.05^62^. The raw matrix was further normalized to TPM (Transcripts Per Kilobase Million) for gene expression quantification. Genes expression data from different tissue was downloaded from a previous publication^55^.

TPM values of genes were clustered by k-means method in the heatmaps in Fig. 5e and 5h. Modules information generated by WGCNA^63^ was used for key gene set selection.

For functional enrichment, GO annotation files were generated from IWGSC Annotation v1.1 and an R package clusterProfiler was used for enrichment analysis^64^.

### Cut&Tag and ATAC-seq Data Processing

Cut&Tag data analysis was largely based on the previously provided pipeline^65^. In brief, two replicates sam files were converted to bam files and sorted by samtools, respectively^59^. We further filter the reads mapped with “samtools view -bS -F 1,804 -f 2 -q 30” to filter the low-quality mapped reads. The high-quality mapped reads were reduplicated using Picard-2.20.5-0 (“Picard Toolkit.” 2019). Two replicate bam files were merged by samtools. For IGV browser visualization, merged bam files were converted to RPKM (Reads Per Kilobase per Million mapped reads) normalized bigwig files with 10 bp bin size for browser visualization by bamCoverage provided by deepTools (3.3.0) with parameters “-bs 10 --effectiveGenomeSize 14,600,000,000 --normalizeUsing RPKM --smoothLength 50”^66^. For Cut&Tag data comparison, scale factors were calculated by ChIPseqSpikeInFree (v1.2.4)^67^. For peak calling, both SEACR v1.3 and MACS2 v 2.1.2 were used^68, 69^. We performed SEACR with numeric threshold 0.05 and normalized stringent model. For narrow histone marker (H3K27ac and H3K4me3) and broad histone markers (H3K27me3), parameters “-p 1e-3” and “--broad --broad-cutoff 0.05” provided by MACS2 were used, respectively. Finally, only peaks generated by MACS2 which overlapped with that generated by SEACR were retained for downstream analysis by “intersect -wa” parameters of bedtools v2.27.1^70^.

The bam file process and bigwig conversion steps in ATAC-seq are the same as that in Cut&Tag. For ATAC-seq peak calling, only MACS2 was used with parameters “--cutoff-analysis --nomodel --shift -100 --extsize 200”. ChIPseqSpikeInFree was not applied in ATAC-seq data process.

For both Cut&Tag and ATAC-seq peaks, if a peak overlapped with the proximal of a gene, including 3kb upstream and 2.5 kb downstream, we assigned the peak to the gene. If multiple genes meet the condition, a position priority strategy (promoter > exon > intron > 5’UTR > 3’UTR > downstream) and nearest gene principle was used for target genes assign. An R package ChIPseeker was used for this peaks annotation process^71^.

### DERs detection

Reads count under special peaks of Cut&Tag and ATAC-seq were calculated by FeatureCount^61^. For Cut&Tag data, scale factors generated by ChIPseqSpikeInFree were used for differential analysis following the suggested usage of DESeq2 method in ChIPseqSpikeInFree manual^67, 72^.

### Chromatin state analysis

For chromatin state analysis, chromHMM was used^40, 41^. “BinarizeBam” and “LearnModel” commands were used for chromatin-state annotation. We used 9 CUT&Tag data as input and got 15 chromatin-states (Fig. 3a left panel).

### ATAC-seq foorprints identity

HINT (Hmm-based IdeNtification of Transcription factor footprints) was used for ATAC-seq footprints identity^73^. JASPAR Plantae database (https://jaspar.genereg.net/) was used as motifs set^74^.

### Distal ACRs annotation

The distal ACRs annotation strategy was largely based on a previous study with a small modification^38^. In brief, genes within 0.5M from a distal ATAC-seq peak are considered candidate target genes (Supplementary Fig. 4b). Then if there is a significant positive correlation between gene expression (TPM) and ATAC-seq signal (FPKM) in eight samples, this gene is considered to be the target of the peak (Supplementary Fig. 4b). A small difference from the Corces et al. (2018) is that we use this strategy only for distal ATAC-seq peaks, and proximal peaks annotation was based on the nearest gene principle^38^.

### GRNs construction in early embryogenesis

GRNs construction in early embryogenesis was based on top-down and bottom-up methods (Fig. S5a). We believe that genes whose expression was up-regulated but not promoter ATAC signal may be largely regulated by TFs that both expression and pACRs up-regulated (Fig. 4b left panel). Thus, the GRN contained four layers, in which the lowe two layers were genes firstly up-regulated in DPA2 and DPA4 compared with DPA0, respectively. Firstly TFs motif was enriched for the up-regulation (DPA2/DPA0) promoter ATAC-seq peaks that were assigned to up-regulated (DPA2/DPA0) genes using homer software (v4.11)^75^. Two types of transcription factors were enriched, including one TCP and two MYB. We used the top-down method to find their downstream target genes through the footprint of the three TFs. Then we used the bottom-up method to find the upstream TFs of genes in the lower two layers through motif enrichment by clusterProfiler^64^. Finally, we got the TFs in the second layer through the intersection of the targets of the top layer and the enrichment result of the bottom layers. These TFs were both expression up-regulated and pACR signal up-regulated. In most conditions, one motif corresponds to multiple TFs. To get the specific TFs, we overllaped our GRN result with previous Genie3 network^55^ and got the final GRN in Fig. 4h. EMB genes were got from a previous review in Arabidopsis^76^, and we used diamond (v2.0.13) with parameter “--id 50 -e 1e-20 -f 6”^77, 78^.

### Psuedotime indexing and GRNs construction in embryo body formation

Only six-module genes generated from WGCNA were used for the analysis in Fig. 6. The psuedotime indexing method is the same as previous studies^79–81^. Briefly, principal component analysis

PCA was used for both RNA- and ATAC-seq data for specific genes. The psuedotime was calculated based on the sample distance between neighbor samples and was scaled to a range from 0 to 10. For one gene, the expression model was fit based on expression level and psuedotime using the “loess” function in R, and 500-time points were generated between 0 to 10, as well as the corresponding expression levels based on the fitted curve.

For one gene, the expression level was normalized by Z-score. We further calculated the PC1 and PC2 for every gene using the expression values of eight samples. Because the standard expression values of all genes can form a circle, the atan2 function in R was used to return the angle in radians for the tangent PC2/PC1, which were further used for gene expression order ranking. As a result, the psuedotime expression of genes was ranked and visualized by complexHeatmap in R^82^. GO and motif (generated from footprint analysis) enrichment was calculated by clusterProfiler^64^.

For GRN building, the TFs for significantly enriched motifs with *p*-value < 0.05 were used. We further calculated the Pearson correlation between TFs and corresponding targets, and only significant pairs (*p*-value < 0.05) were retained. To simplify the GRN, we focused on the TFs-TFs network. As a result, several TFs contained multiple genes, we combine those as TF modules, including TCPs, ARFs, MYBs, and woxs.

## Data availability

The raw sequence data were deposited in the Genome Sequence Archive (https://bigd.big.ac.cn/gsa) under accession number xxx.

## Acknowledgements

We thank Professor Falong Lu and Zhuo Du from IGDB-CAS for critical comments on the manuscript. This research was supported by the Strategic Priority Research Program of the Chinese Academy of Sciences (XDA24010204) and the National Natural Sciences Foundation of China (31970631) to J.X.

## Author contributions

J.X. and L.Z. designed and supervised the research and wrote the manuscript. H.Z., L.Z. and X.-L.L. performed Cut&Tag, ATAC-seq and RNA-seq experiments; L.Z. performed data analysis; X.-L.L., Y.-M.Y and H.W. performed the rest of the experiments. All authors discussed the results and commented on the manuscript.

---

**Fig. S1.**
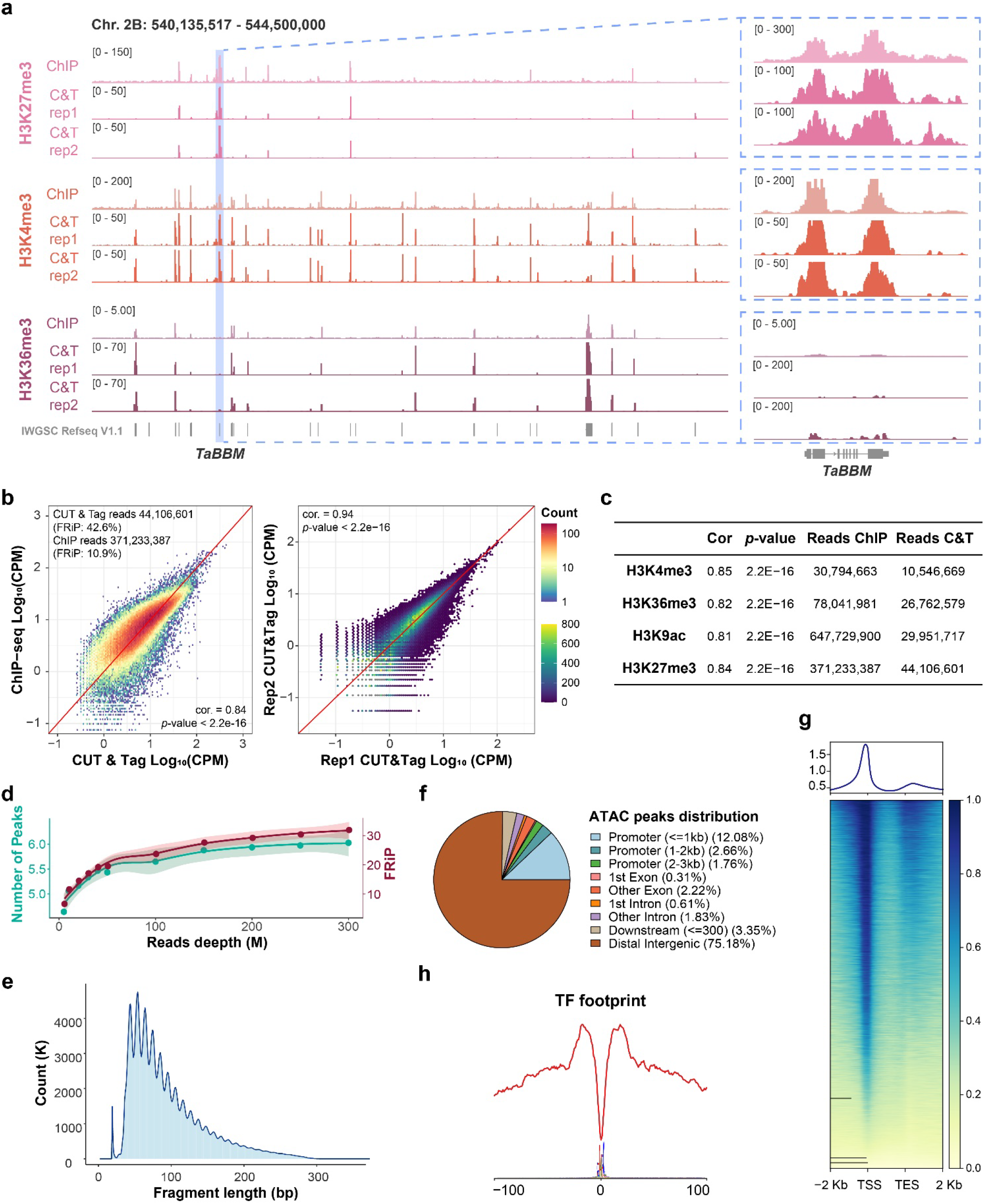
Quality control for CUT & Tag and ATAC-seq. **a-c**, CUT & Tag recapture ChIPseq signal with high reproducibility and low sequencing depth. IGV (Integrated Genome Browser) showing high reproducibility between ChIP-seq and CUT & Tag in seedlings (**a**). Correlation between ChIP-seq and CUT & Tag as well as two biological replicates (**b**). Sequencing depth comparison for various histone modifications between CUT&Tag and ChIPseq (**c**). **d-h**, Quality control of ATAC-seq data. Peaks number, and FRiP (The fraction of reads in called peak regions) were calculated at different sequence depths (**d**). Fragment size distribution (**e**) and peak distribution within the genome (**f**). ATAC-seq signal distribution along with genes (**g**). Transcription factor footprint profile of ATAC-seq data (**h**).

**Fig. S2.**
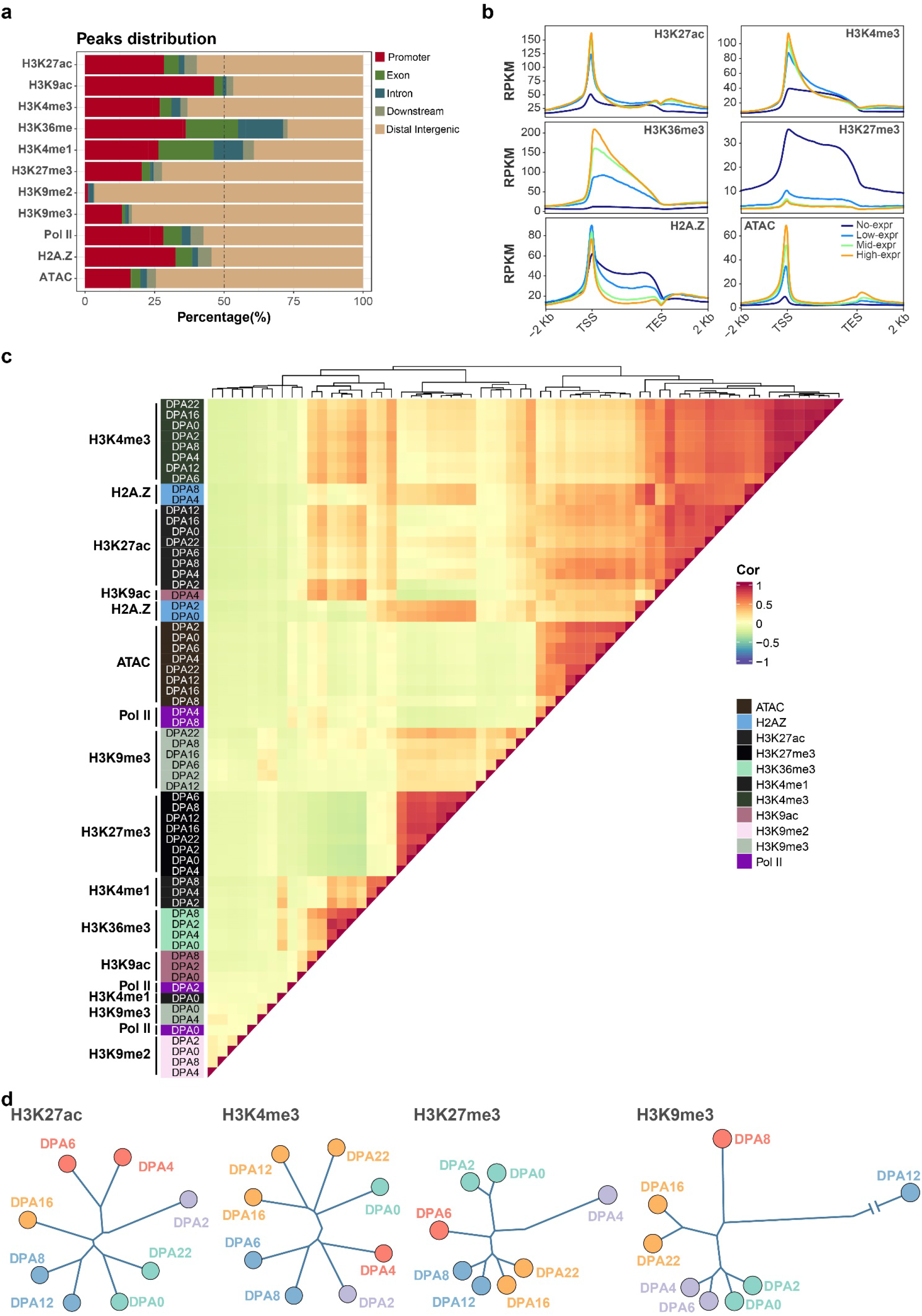
Features of various histone modifications. **a**, Peak distribution of different histone modifications relative to genes. **b**, Correlation between different types of histone modification profiles and gene expression levels. **c**, Pair-wise correlation map among different histone modification profiles. Jaccard index was calculated based on the peaks overlap, and then Pearson correlation scores were generated. **d,** Cluster dendrogram of histone modifications. Reads count in peaks were normalized by CPM, and euclidean distances were calculated for tree clustering.

**Fig. S3.**
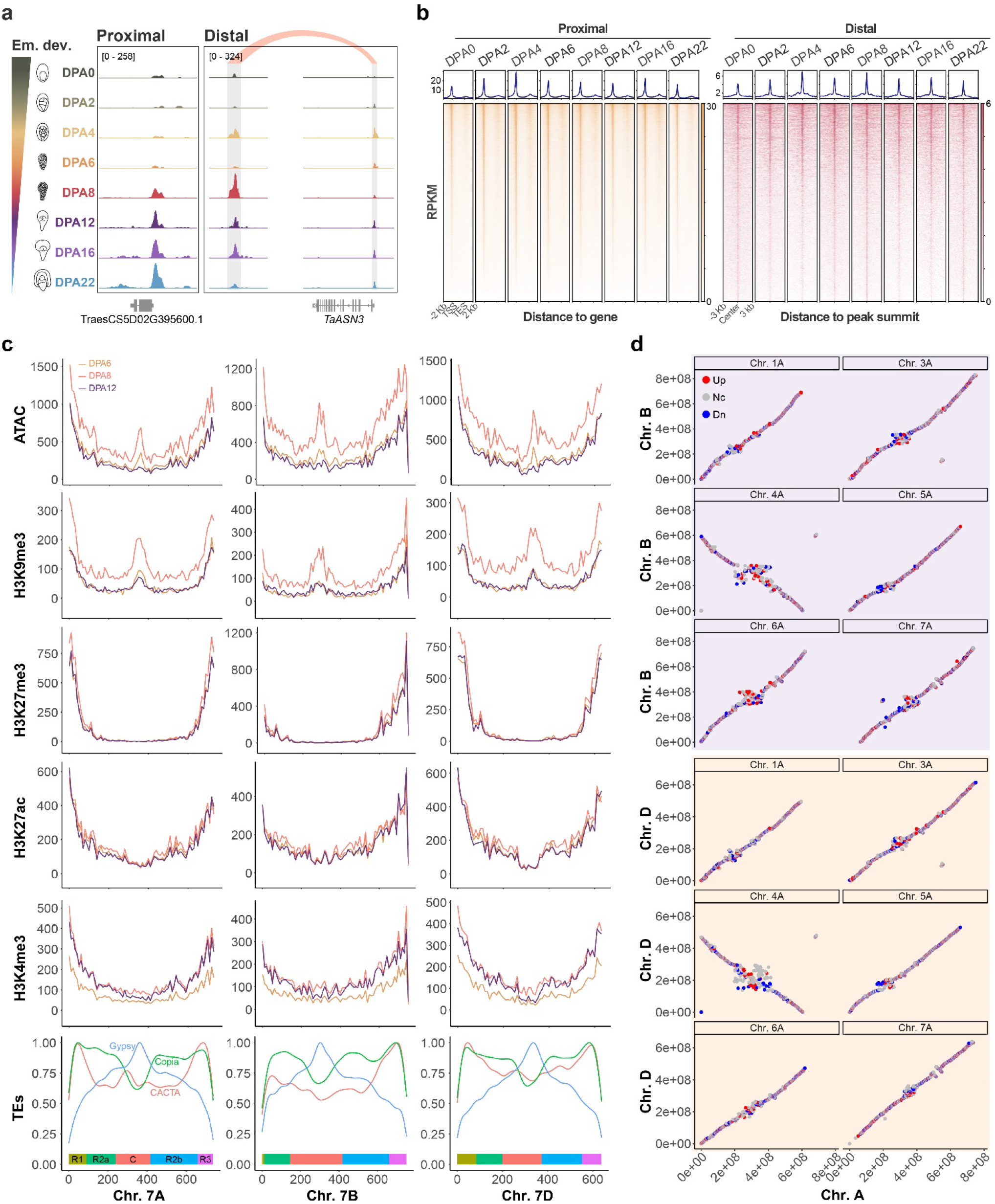
Dynamic ACRs during wheat embryogenesis and difference among sub-genomes. **a**, IGV showing chromatin accessibility change cross wheat embryogenesis in both proximal and distal regions. **b**, ACRs dynamic in proximal (left) and distal regions (right) during wheat embryogenesis. **c**, Chromatin accessibility (ATAC), histone modifications and various types of TEs distributions change on Chr. 7 of A-, B- and D-subgenome at DPA 6, 8 and 12. Color bars on x and y axis indicate the chromosome segments defined by Consortium (IWGSC) et al., 2018. **d**, Comparisons of chromatin accessibility of collinear regions among subgenomes. Down- (Dn), and Up-regulated (Up) loci were define by the log_2_(fold change Chr. B/Chr. A) (left top-panel) and log_2_(fold change Chr. D/Chr. A) (left bottom-panel) values bigger and lower than 1 and -1. Nc indicates no change between subgenomes.

**Fig. S4.**
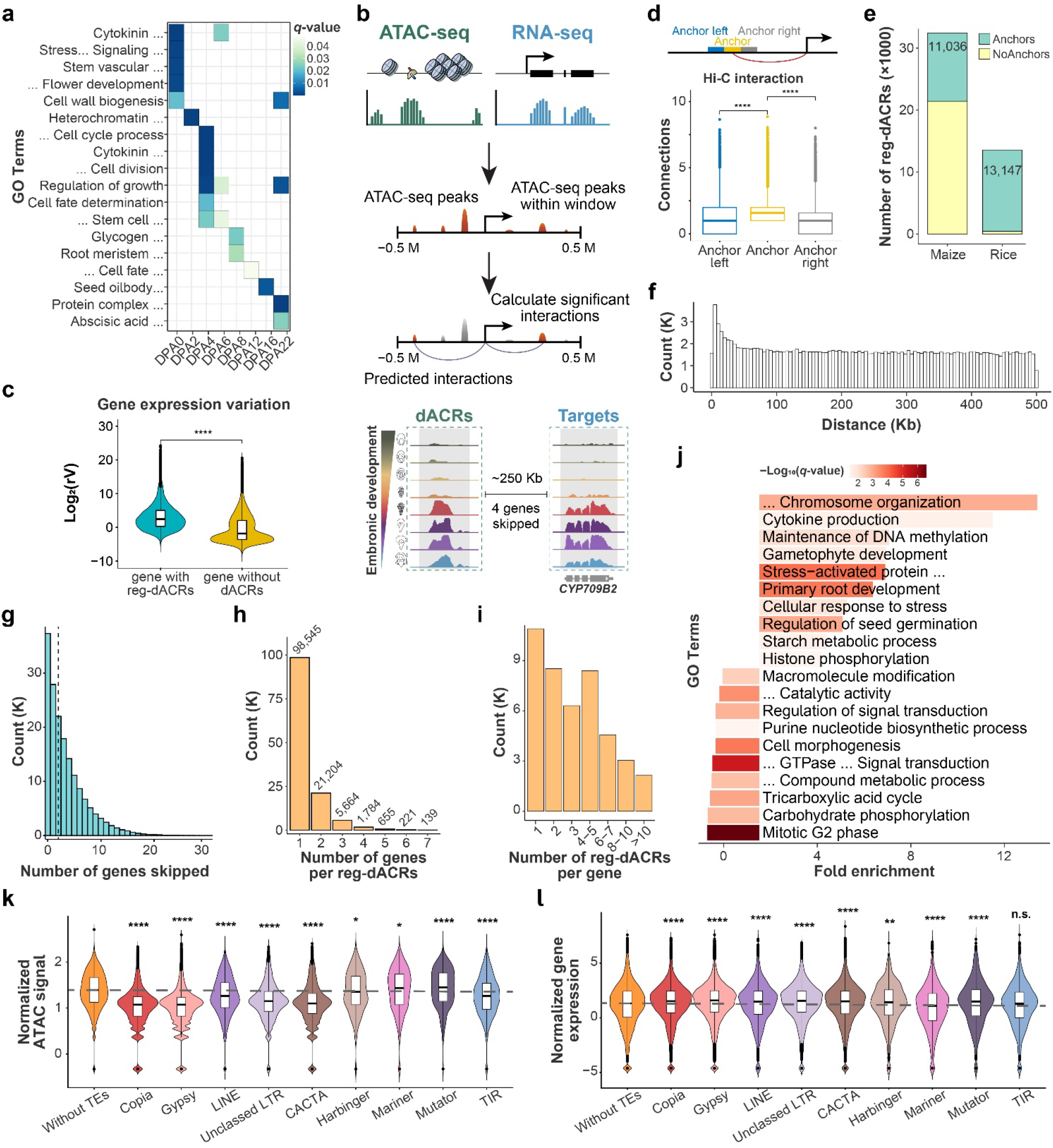
Association between reg-dACRs to genes. **a**, GO enrichment of stage-specific genes with pACRs. **b**, Schematic of the strategy used to link distal ATAC-seq peaks to genes (see Method for detail). **c**, Relative variation of expression level for genes with dACRs or without. **d**, Hi-C anchors highlight the connection between reg-dACRs and target genes. **e**, Number of dACRs homologous segment in maize and rice. Based on previously published Hi-C and ChIA-pet data, the homologous segment was grouped in anchors and NoAnchors. **f, g,** Distance of reg-dACR to genes (**f**) and the number of genes skipped (**g**) between reg-dACR and target genes. **h**, Number of genes can be targeted per reg-dACRs. **i,** Number of reg-dACRs distribution for the individual target gene. **j,** Corresponding GO enrichment for genes with solo or multiple dACRs. **k**, **l**, Comparison of ATAC-seq signal and corresponding target genes expression among reg-dACRs overlapped by different types of TEs. Mann-Whitney U test (two-sided) was used for **c** and **k** (* : p <= 0.05; ** : p <= 0.01; ***: p <= 0.001).

**Fig. S5.**
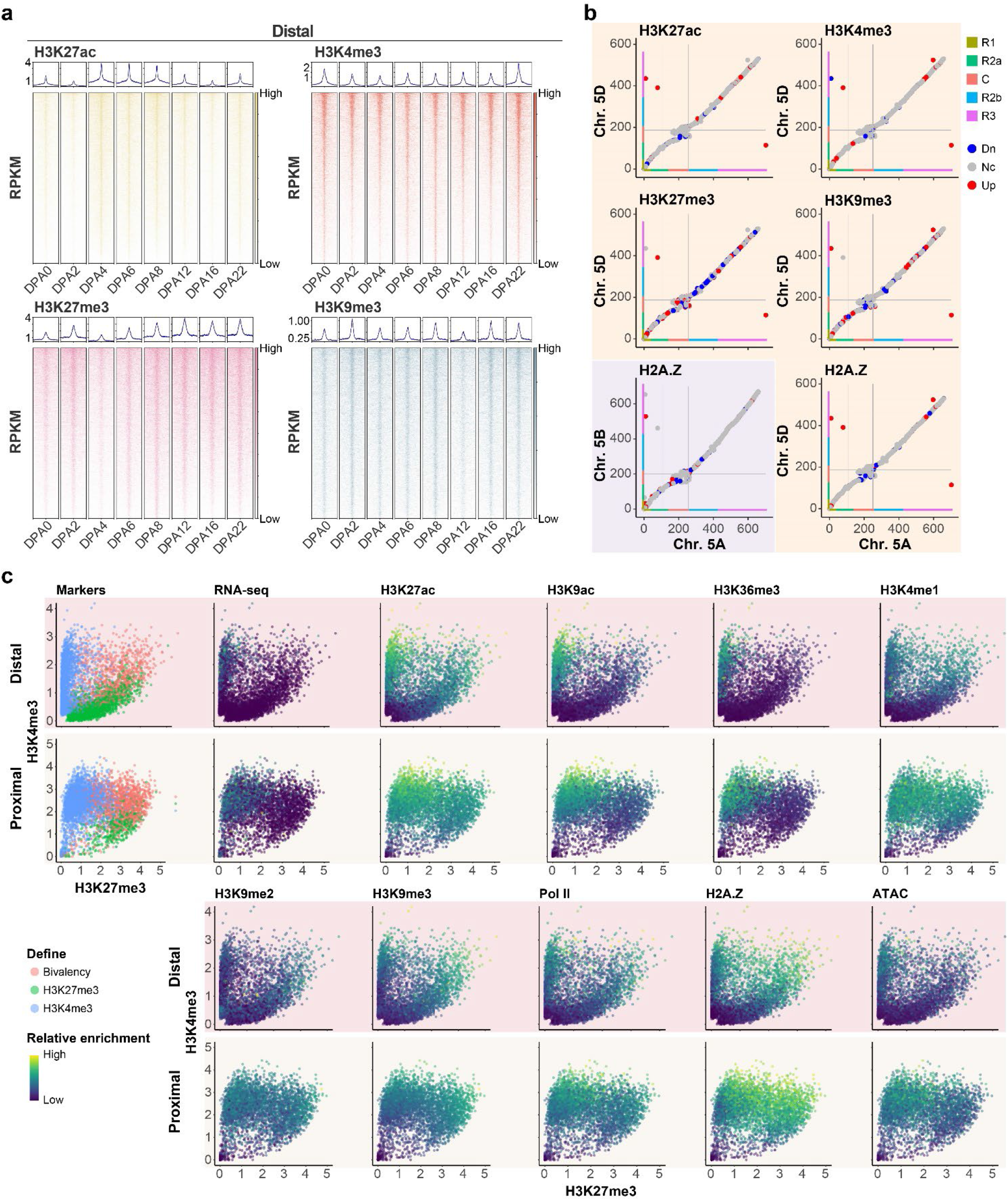
Histone modifications dynamic and subgenome comparison. **a,** Histone modifications dynamic in distal regions during wheat embryogenesis. **b**, Comparisons of histone modifications H3K27ac, H3K4me3, H3K27me3, H3K9me3 and H2A.Z of collinear regions between three subgenomes of Chr. 5. Down- (Dn), and Up-regulated (Up) loci were defined by the log_2_(fold change Chr. 5B or Chr. 5D / Chr. 5A) values bigger and lower than 1 and -1, respectively. Nc indicates no change. Color bars and grey lines indicate the chromosome segments and centromere regions as those in Fig. 2d. **c**, Scatter plots correlating H3K27me3 (x axis) and H3K4me3 (y axis) enrichments (log_2_CPM) at distal and proximal regions. Color represents chromatin state defined as indications, gene expression and multiple histone modifications level (log_2_CPM).

**Fig. S6.**
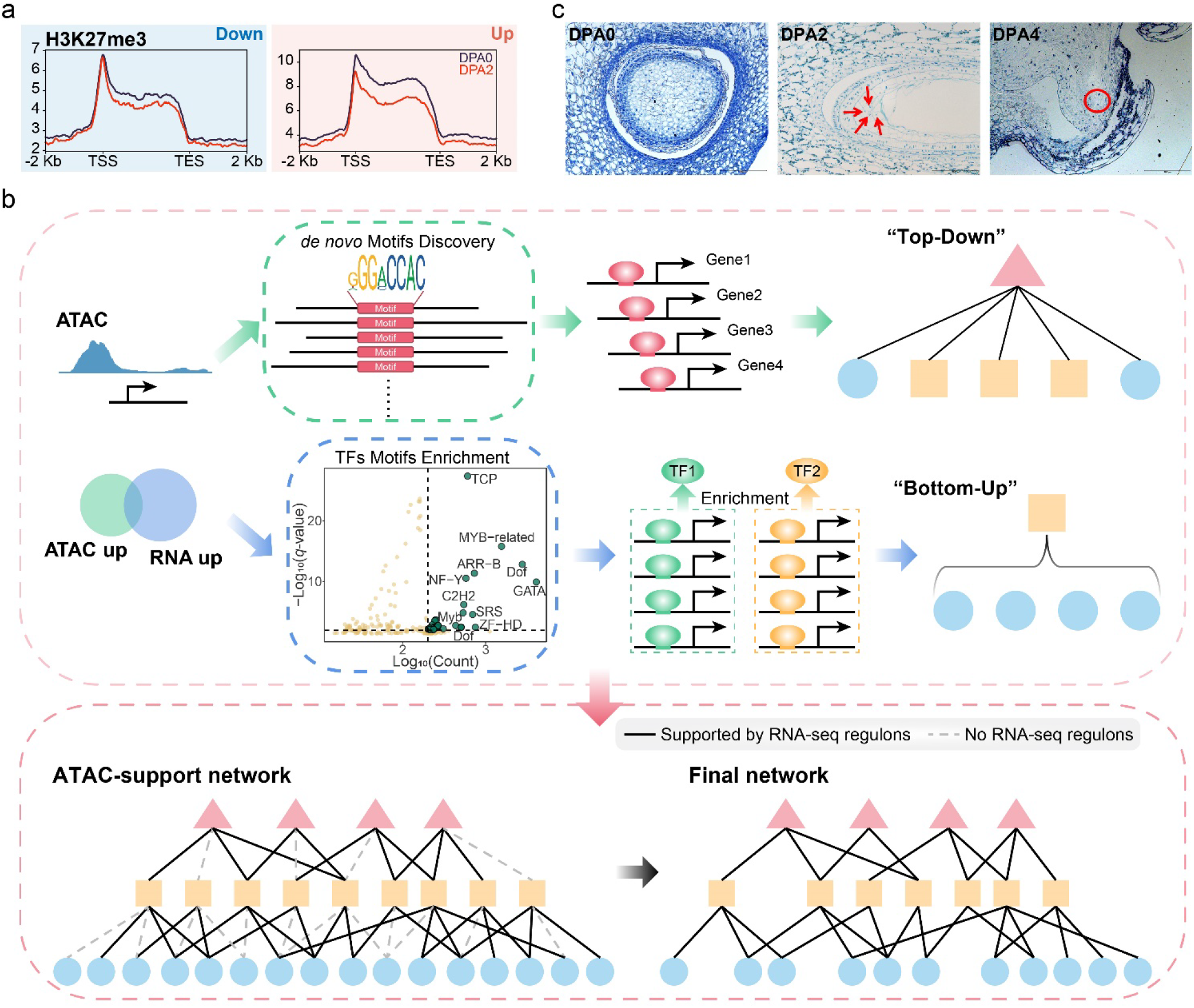
Histone modification change and build-up of transcription regulation network for ZGA. **a**, H3K27me3 change following fertilization at dn- and up-regulation gene sets as Fig. 4b. **b**, Schematic of the strategy for GRNs building (see method for detail). **c**, Morphological change of embryo-sac during early embryogenesis.

**Fig. S7.**
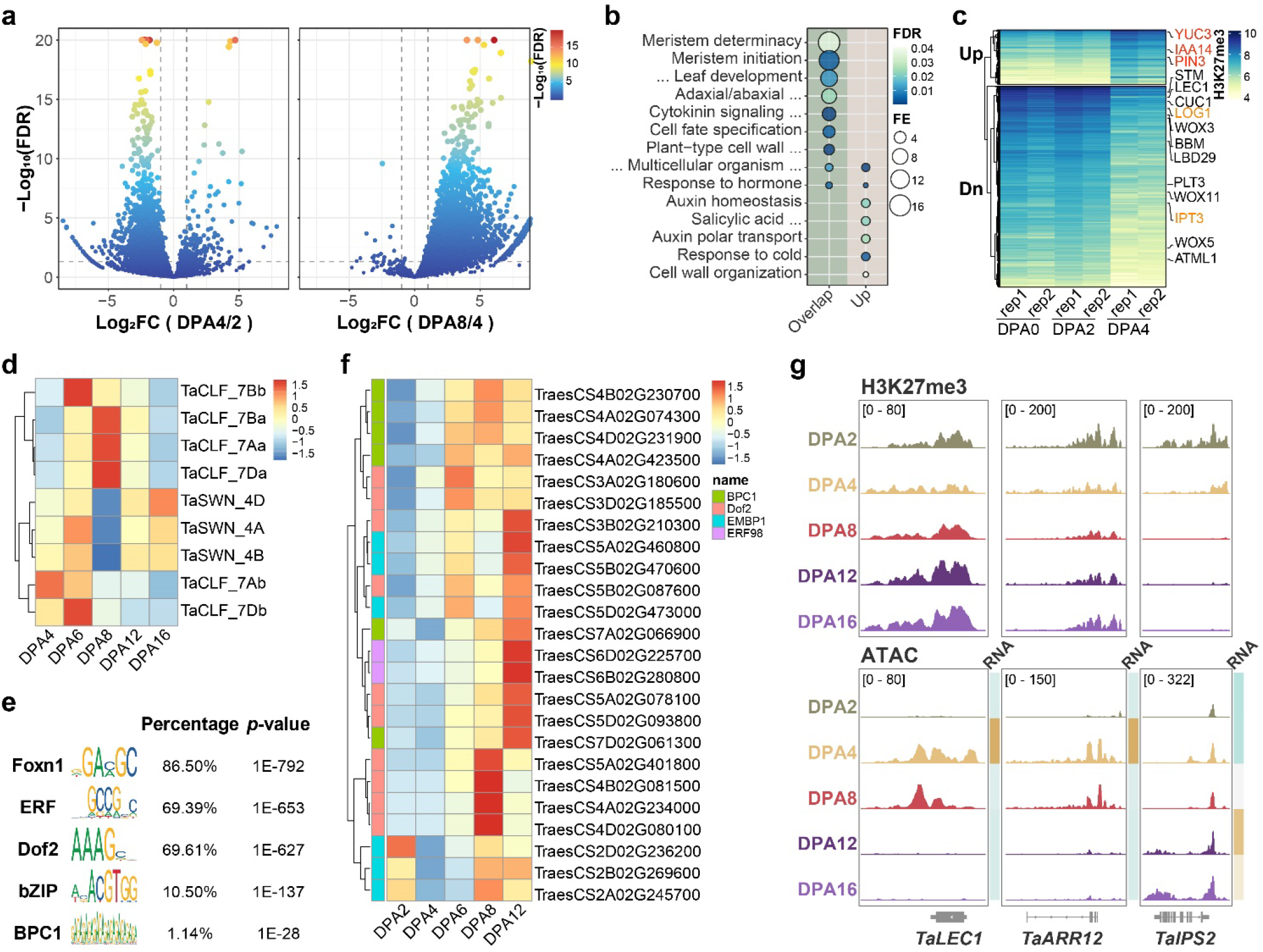
H3K27me3 reprogramming at pre-embryo. **a**, Volcano plot of differential H3K27me3 enriched regions between DPA2 and DPA4 (left), and between DPA4 and DPA8 (right). Differential regions were defined by the threshold absolute value of log_2_(Fold Change) ≥1 and FDR ≤0.05 by DESeq2. **b,** GO enrichment of gene sets which lost H3K27me3 at DPA4 and re-gain at DPA8, and gene sets which *de novo* gain of H3K27me3 at DPA8. **c**, Heatmap showing differential H3K27me3 modification genes between DAP4 and DAP2. Embryonic development essential genes were highlighted. Auxin-related genes were labeled in red and CK-related genes were labeled in orange. **d**, The expression level of H3K27me3 methyltransferase coding genes in selected embryonic development stages. **e**, **f**, Motif enrichment of H3K27me3 deposition loci at DAP8 (**e**) and the expression levels of cognate binding TFs (**f**). Homer was used for motif enrichment analysis. **g**, H3K27me3 and chromatin accessibility dynamics at embryo essential genes *TaLEC1, TaARR12* and *TaIPS2*.

**Fig. S8.**
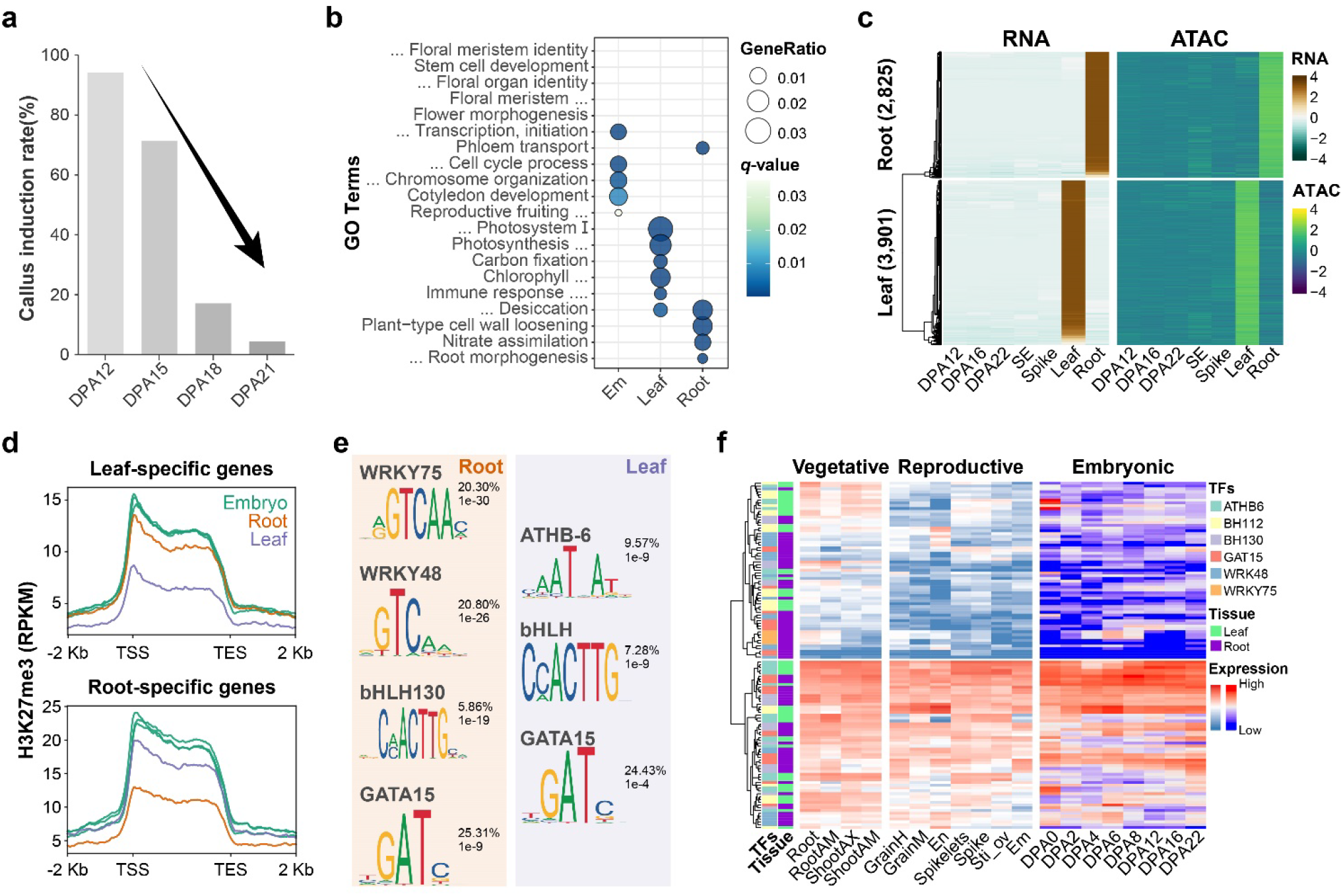
Inhibition of deep organogenesis in late embryo at chromatin level. **a**, Callus induction rate drops as embryo developing after DPA 12. **b**, GO enrichment of tissue identity genes found in Fig. 7d. **c,** Synchronous patterns between root- and leaf-identity genes expression and corresponding pACRs in individual tissue. **d,** H3K27me3 modification levels on root- and leaf-specific genes in individual tissue. **e**, Motifs enrichment within regulatory regions of root- (left) and leaf-specific genes (right). Percentage showed the presence frequency, while P-value indicates the enrichment. **f**, Expression patterns of candidate TFs that can bind motifs found in **e**, cross different tissues and during embryogenesis.

**Fig. S9.**
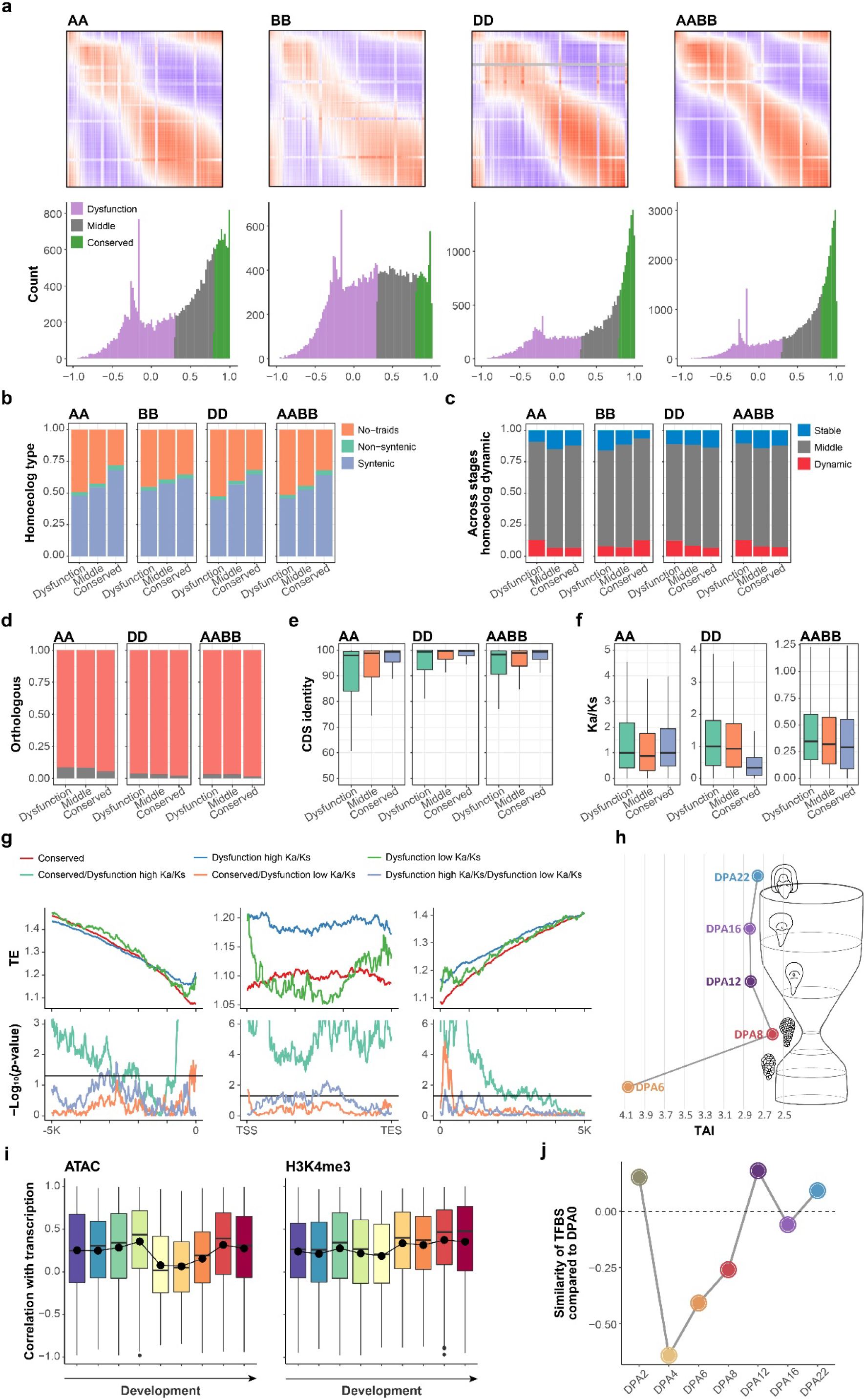
Comparisons of embryogenesis among different ploidy wheat and hourglass model. **a**, Comparisons of gene expressions between hexaploid wheat (AABBDD) and ancestors (AA, BB, DD and AABBDD) and corresponding Pearson correlation index. Each row represents an individual gene in hexaploid wheat and each line represents an individual gene in ancestor. Pearson correlation index was from -1 to 1, and indicated by colors from blue to red. Genes were clustered into three categories based on Pearson index from the comparisons between hexaploid wheat and ancestors, which are dysfunction, middle and conserved (bottom). **b**, **c,** subgenome synteny (**b**) and triad dynamic expression across developmental stages (**c**) for gene sets defined in **a**. **d**, Percentage of genes has paralogous in corresponding ancestors for gene sets defined in **a**.. **e**, Sequence similarity between hexaploid wheat and different ancestors for the corresponding gene sets defined in **a**. **f**, Ka/Ks comparisons for different gene sets defined in **a**. **g**, TEs insertion frequency at different positions relative to different gene sets (top) and comparisons between different gene sets (bottom). Mann-Whitney U test (two-sided) was used for the significant difference. **h**, TAI index for expressed genes from DPA6 to DPA22, which represent an hourglass model. **i,** ATAC but not H3K4me3 contributed to genes expression across wheat embryogenesis. **j**, Similarity of TFs binding pattern compared with that in DPA0. All TFs binding patterns were calculated by chromVAR.

**Fig. S10.**
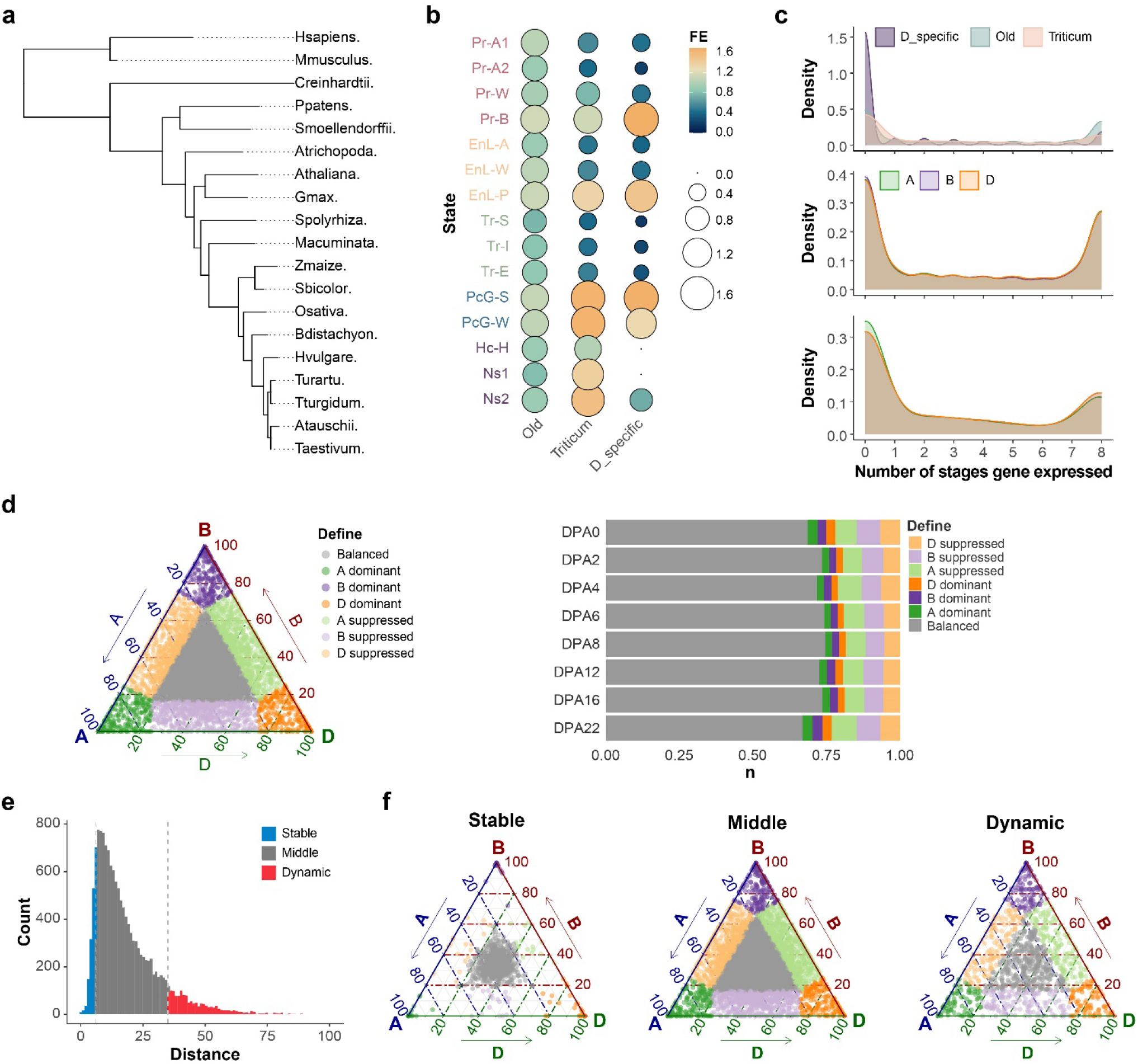
Chromatin landscape affected sub-genome bias expression. **a**, Phylostratigraphic map of various wheat species **b**, Chromatin states enrichment for different evolutionary age genes in wheat embryo. **c**, Expression spectrum of different evolutionary age genes in wheat embryo. **d**, Define of homeologs bias expression (left) and percentage of different homeologs bias expression types across eight wheat embryonic developmental stages (right). **e**, Homeologs dynamic expression across wheat embryogenesis. Euclidean distance method was used to analysis as (Ramírez-González et al., 2018). **f**, Homeologs bias expression for gene sets defined in **e**.

**Fig. S11.**
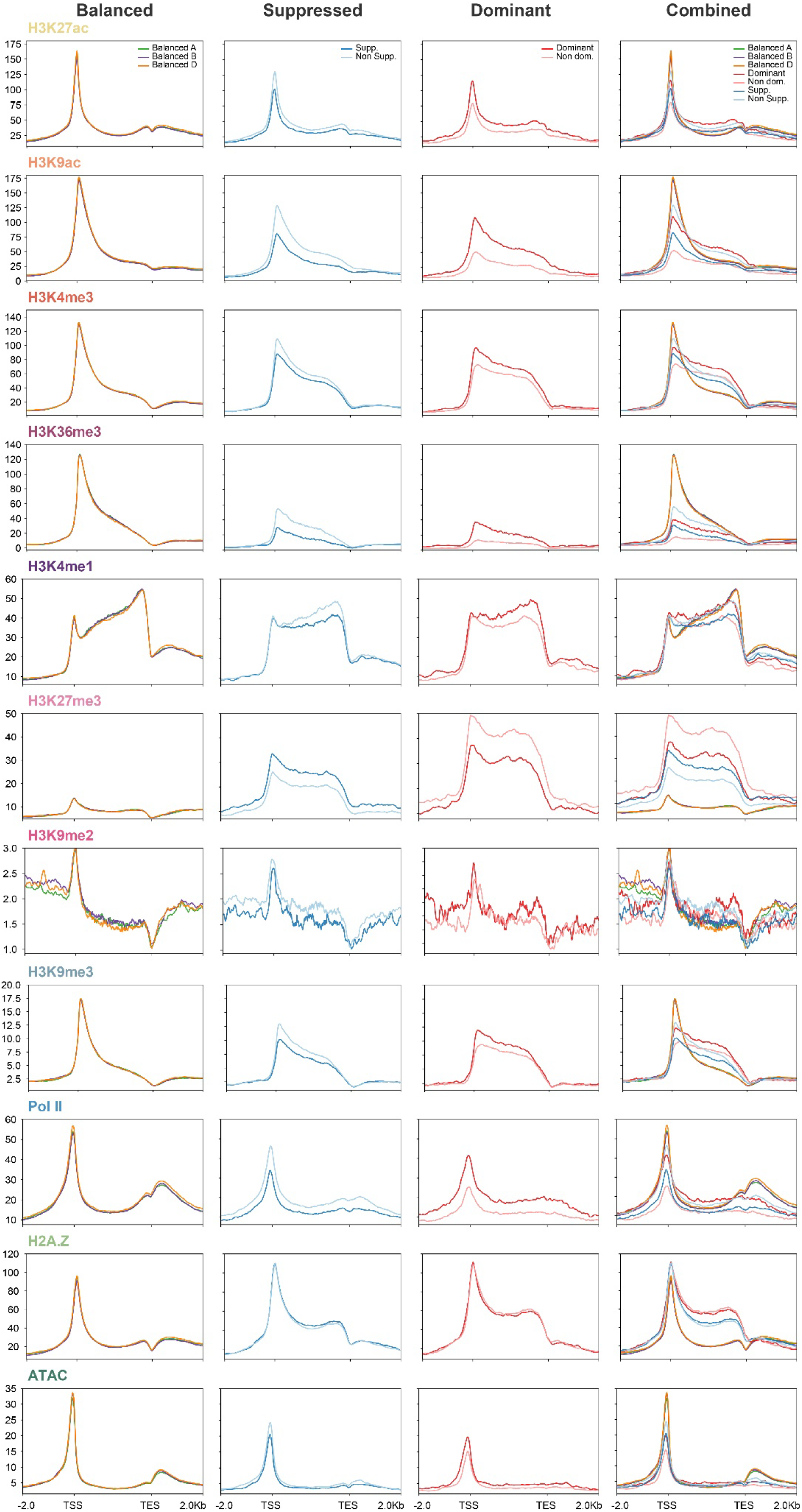
Epigenetic modification on different types of homeologs bias expressed genes. Balanced, suppressed, and dominant genes were shown in the first three columns, and a combination of different types of homeologs was shown in the fourth column. Epigenetic modification signals were normalized using RPKM, with 10 bp bin size.

**Fig. S12.**
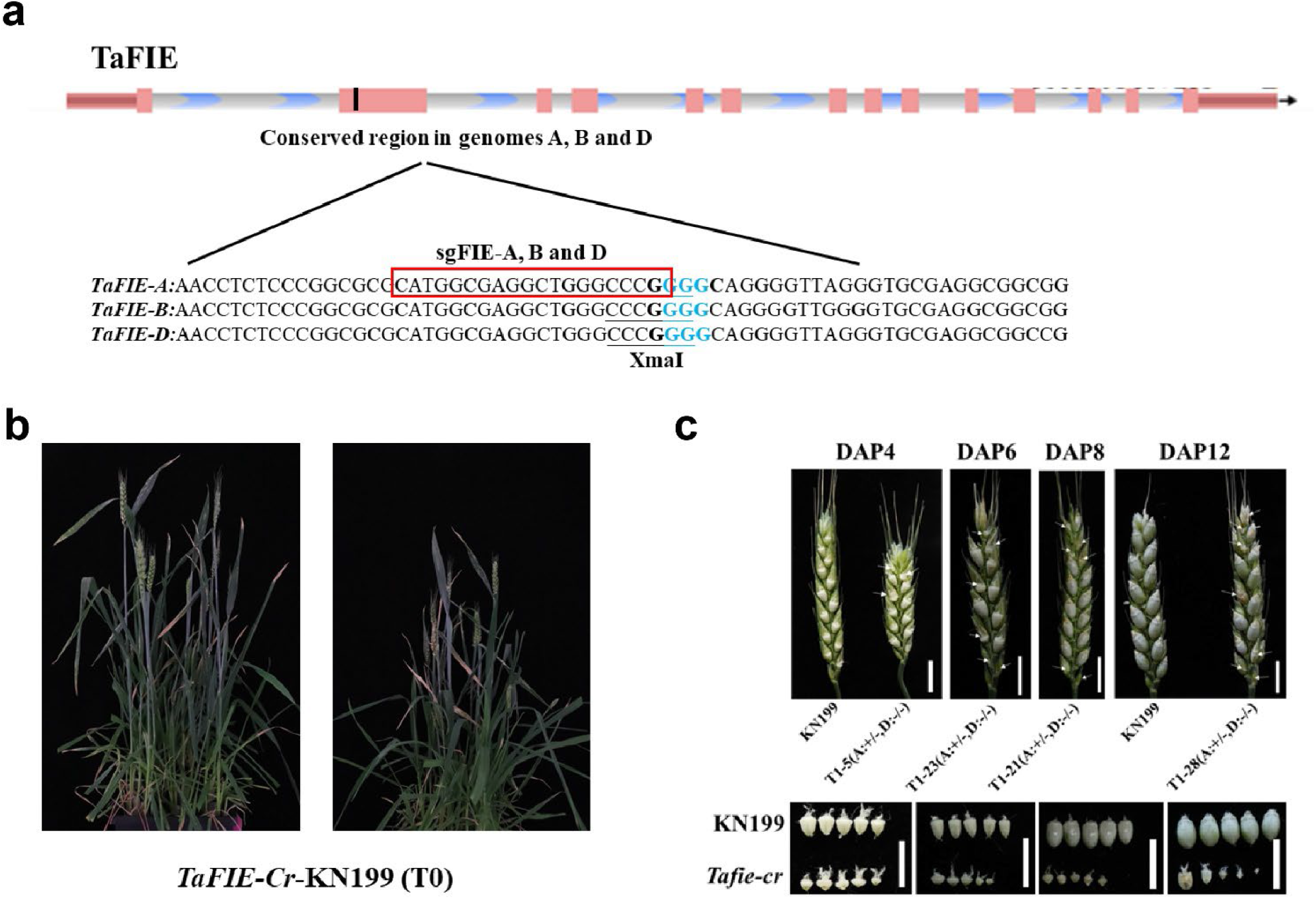
Loss of core component of PRC2 led to aberrant seeds in wheat. **a**, Genotyping of CRISPR plant at *TaFIE*. **b**, Growth phenotype of *TaFIE-Cr-KN199* as compared to wildtype KN199. **c**, Aberrant seed produced by *TaFIE-Cr-KN199*.

